# Placebo analgesia does not reduce empathy for naturalistic depictions of others’ pain in a somatosensory specific way

**DOI:** 10.1101/2020.12.29.424686

**Authors:** Helena Hartmann, Federica Riva, Markus Rütgen, Claus Lamm

**Author notes:** Corresponding author, Social, Cognitive and Affective Neuroscience Unit, University of Vienna, Liebiggasse 5, 1010 Vienna, Austria.

## Abstract

Empathy for pain involves the affective-motivational and sensory-discriminative pain network. The shared representations account postulates that sharing another’s pain recruits underlying brain functions also engaged during first-hand pain. Critically, causal evidence for this has only been shown for affective pain processing, while the specific contribution of one’s own somatosensory system to empathy remains controversial. Experimental paradigms used in previous studies did not a) direct attention towards a specific body part or b) employed naturalistic depictions of others’ pain, which could explain the absence of somatosensory effects. In this preregistered fMRI study, we thus aimed to test whether a causal manipulation of first-hand pain affects processing of empathy in a somatotopically-matched manner. Forty-five participants underwent a placebo analgesia induction in the right hand and observed pictures of right vs. left hands in pain. We found neither behavioral nor neural evidence for laterality-specific modulation of empathy for pain. However, exploratory analyses revealed a general effect of the placebo on empathy, and higher brain activity in bilateral anterior insula when viewing others’ hands in pain corresponding to one’s own placebo hand. These results refine our knowledge regarding the mechanisms underlying empathy for pain by specifying the influence of first-hand pain on empathic responding.

## 1 Introduction

Neuroimaging studies of empathy for pain have consistently found involvement of a network of brain regions related to affective-motivational processing of pain, such as anterior midcingulate cortices (aMCC) and anterior insula (AI). Other studies have highlighted the importance of sensory-discriminative processing in empathy, e.g., in primary (S1) and secondary (S2) somatosensory cortices (Keysers et al., 2010 for a review; Fallon et al., 2020; Fan et al., 2011; Jauniaux et al., 2019; Timmers et al., 2018 for meta-analyses). However, the exact role of the latter component during empathy remains controversially discussed. More specifically, it is unclear whether somatosensory pain processing networks contribute to empathy for pain by means of somatotopic matching, as they do for first-hand pain, for example, whether we recruit somatosensory representations of our right hand, when empathizing with the right hand of another person in pain. Resolving this issue was the main aim of the present study.

Empathy constitutes a complex, multi-faceted process involving cognitive, affective, and behavioral mechanisms (see e.g., Marsh, 2018; Weisz & Zaki, 2018 for recent reviews). Here we refer to empathy as an affective state isomorphic to the affective state of another person that includes a partial and experiential sharing of the other’s affective state (de Vignemont & Singer, 2006; Hall & Schwartz, 2019; Lamm et al., 2019; Shamay-Tsoory & Lamm, 2018). Interestingly, affective and somatosensory brain regions involved in processing empathic pain overlap in part with brain regions that are also active when we experience pain ourselves (Lamm et al., 2011; Singer et al., 2004). Within the so-called shared representations account of empathy, this activation overlap has been taken to suggest that shared representations may underlie such vicarious experiences, in other words, that we might recruit the same neural substrates when feeling pain ourselves and when empathizing with someone else in pain (Bastiaansen et al., 2009; Bernhardt & Singer, 2012). Although the majority of studies report this overlap of activation in aMCC and AI (Benuzzi et al., 2018; Corradi-Dell’Acqua et al., 2011; Jackson et al., 2006), other studies have pointed to S1 and/or S2 as equally relevant for pain empathy (Avenanti et al., 2005; Bufalari et al., 2007; Fabi & Leuthold, 2017; Gallo et al., 2018; Lamm, Nusbaum, et al., 2007; Motoyama et al., 2017; Novembre et al., 2015; Riečanský & Lamm, 2019 for a review). Evidence for shared representations was recently also confirmed on the level of multivariate patterns, pinpointing the mid- to anterior insular cortex as an area in which self/other pain representations are shared (Zhou et al., 2020; but see also Krishnan et al., 2016).

Importantly, an overlap in activation, as demonstrated e.g. with fMRI, does not necessarily imply shared underlying functions and has fostered an active debate regarding the existence and location of shared representations underlying first-hand and empathy for pain. Trying to go beyond correlational evidence of shared activations towards evidence for shared representations, recent studies have made use of causal and/or psychopharmacological manipulations (Gallo et al., 2018; Rütgen et al., 2018; Rütgen, Seidel, Riečanský, et al., 2015; Rütgen, Seidel, Silani, et al., 2015; Rütgen et al., 2020). One such method is placebo analgesia, the reduction of first-hand pain by means of an inert pharmacological substance, which has been found to decrease both first-hand pain ratings and pain-related brain activity in regions such as insula, ACC, S1, S2 and thalamus (Atlas & Wager, 2014 for a metaanalysis; Eippert et al., 2009; Geuter et al., 2013; Wager et al., 2004; see Colloca et al., 2013; Wager & Atlas, 2015 for reviews). Moreover, placebo analgesia has been successfully employed to downregulate pain via localized manipulations, e.g., using gels or creams on different body parts (Bingel et al., 2006; Geuter et al., 2013; Schafer et al., 2015; Schenk et al., 2014; Wager et al., 2004). Turning to empathy, Rütgen and colleagues used this method in three consecutive studies to test whether such a first-hand pain reduction transfers to empathy for pain, as the shared representations account would suggest (Rütgen et al., 2018; Rütgen, Seidel, Riečanský, et al., 2015; Rütgen, Seidel, Silani, et al., 2015). Indeed, global placebo analgesia by means of a placebo pill lowered activity in brain regions such as AI, dorsal ACC, S2 and thalamus, thus demonstrating that this induction procedure was indeed able to down-regulate first-hand somatosensory processing (Rütgen, Seidel, Silani, et al., 2015). In the empathy condition, the authors further observed lower ratings of unpleasantness and empathy for pain as well as lower activity in aMCC and left AI in the placebo group compared to a control group who had not received any pill. They further showed a crucial role of the opioid system, as blocking placebo analgesia using the opioid antagonist naltrexone blocked the manipulation’s effects on first-hand and empathy for pain. The behavioral results have since been replicated by an independent research group using real painkillers (Mischkowski et al., 2016) and extended to empathy for positive emotional states (Mischkowski et al., 2019). Similarly, hypnotic analgesia decreased brain responses of both self- and other-related pain in right AI and amygdala (Braboszcz et al., 2017).

Notwithstanding the important advances made by this line of research, several points are still unclear: First of all, these studies did not find any placebo-related modulation of the sensory-discriminative component of pain during empathy, suggesting that empathy for pain might be specifically focused on sharing the affective representation of another’s pain (Rütgen, Seidel, Silani, et al., 2015). Relatedly, it has been proposed that specific types of empathy for pain paradigms recruit somatosensory brain regions to a larger degree, namely tasks where the attention is directed to the somatic location of pain, such as the exact body part in which the pain is inflicted (Lamm et al., 2011; Xiang et al., 2018; Zaki et al., 2016). Note in this context that two types of paradigms are commonly used in pain empathy research: *cue-based* tasks, which use abstract cues to indicate experimentally induced painful (electrical or thermal) stimulation in different intensities delivered to another person (usually on the hands or arms); *picture-based* tasks, which show pictures or pre-recorded videos of individuals’ body parts or faces in pain. Crucially, cue-based tasks usually employ an additional first-hand pain condition where the participants receive the same stimulation they are empathizing with (either in a separate block, or intermixed with the empathy condition), while picture-based tasks focus mostly on the empathic reaction to visual stimulus material (but see Krishnan et al., 2016). One main difference between these paradigms is thus the way one infers the pain of another person. In cue-based tasks, participants rely on implicit cues representing the target or intensity of the stimulation, while picture-based tasks show explicit and directly visible pain events. Importantly, previous studies on placebo analgesia did not use a setup directing participant’s attention to the location of pain, which could possibly have led to reduced sensitivity in observing somatosensory modulation (Rütgen, Seidel, Riečanský, et al., 2015; Rütgen, Seidel, Silani, et al., 2015). Most studies in the past employing picture-based tasks have not employed causal manipulations such as placebo analgesia. Only one line of research used transcranial magnetic stimulation to measure changes in corticospinal motor representations of hand muscles in individuals observing hands or feet being penetrated by needles (e.g., Avenanti et al., 2005; see Keysers et al., 2010 for a review). They showed a reduced amplitude of motor-evoked potentials in the muscles that participants saw being pricked by the needle, which correlated with subjective ratings of sensory pain qualities. This again highlights the importance of somatic resonance and a direct, bodily matching of specific sensory aspects of others’ pain.

We thus conducted a project employing a localized placebo analgesia manipulation on the right hand only (with the left hand acting as a control) and two paradigms tailored to investigate somatosensory involvement in empathy, making it possible to investigate shared representations related to somatotopic, i.e. location-specific, matching. The first part of this project aimed to refine the results of Rütgen, Seidel, Silani, et al. (2015) by using an adapted version of their task, where electrical stimulation was administered either to the participants or a confederate (Hartmann et al., 2021). In this setup, however, while increasing the focus on the affected body part, pain still had to be inferred via abstract cues indicating how painful stimulation of that body part was. In brief, this task and part of the study did not reveal evidence for a location-specific somatosensory sharing of others’ pain. The second part of this project, reported here, aimed to investigate whether or not these findings also hold when using a picture-based task consisting of naturalistic depictions of others in everyday painful situations. The goals of the present, preregistered study were therefore to a) investigate whether placebo analgesia modulates the behavioral and neural responses to explicit naturalistic depictions of others in everyday painful situations, and b) to resolve whether somatosensory brain areas are modulated in a location-specific manner in such a setup. We hypothesized that behavioral and neural responses (both in affective and somatosensory brain regions) would be lower for stimuli corresponding to the right hand, where placebo analgesia was induced, compared to left hand-related stimuli. Our preregistered main predictions were, therefore, that neural somatosensory representations engaged by firsthand pain are also recruited when empathizing with the pain of another person, and that the hand-specific lateralized reduction of first-hand pain would result in a corresponding lateralized reduction of empathy for pain.

## 2 Materials and Methods

### 2.1 Data and code availability statement

Unthresholded statistical maps are available on NeuroVault (https://neurovault.org/collections/9244/). The stimuli of the picture-based empathy for pain task are available and will be shared individually upon request.

### 2.2 Preregistration

In line with the suggestion for openly stating transparency by Simmons et al. (2012), we report how we determined our sample size, all data exclusions, all manipulations, and all measures in the study. This study was preregistered on the OSF prior to any creation of data (Hartmann et al., 2018; preregistration: osf.io/uwzb5; addendum: osf.io/h7v9p). The here reported part of the project was designed to extend the results of Rütgen, Seidel, Silani, et al. (2015) regarding a transfer of first-hand placebo analgesia to empathy for everyday painful situations. The study design and procedures reported here are therefore largely identical to the ones employed in Hartmann et al. (2021), and reproduced here and in the Supplement. In the following methods and results, we clearly separate preregistered procedures and analyses from those added post hoc. Preregistered tests are additionally marked with a ‘p’ in all results-related figures.

### 2.3 Participants

For estimating the sample size, we conducted an *a priori* power analysis using GPower (Faul et al., 2007), using a conservative average of the lowest effect sizes from previous studies (one-sided paired *t*-test; see Rütgen, Seidel, Riecansky, & Lamm, 2015; Rütgen, Seidel, Silani, et al., 2015). We aimed to detect a medium effect size of Cohen’s *d =* .40 at the standard .05 α error probability with a power of 1-ß = 0.8, yielding a sample size of 41 participants. However, considering that the modulation of placebo analgesia might not be equal for empathy in everyday painful situations, 45 placebo responders was set as the data collection stopping-rule. Due to the nature of our research question, it was crucial to obtain a final set of participants who showed a first-hand placebo analgesia effect, in order to investigate a transfer of this effect to empathy for pain. We thus evaluated non-responders to the placebo analgesia manipulation using a set of criteria consisting of three measures already employed in Rütgen, Seidel, Silani, et al. (2015) and a fourth additional criterion possible due to our within-subjects design. In brief, those criteria were 1) strong doubts about the cover story, 2) low belief in the effectiveness of the “medication”, 3) a high number of conditioning trials, and 4) higher first-hand pain ratings for the right/placebo compared to the left/control hand in another task (see also sections M.1 in the Supplement for screening and exclusion criteria, and M.2 for nonresponder identification). From a total of 78 recruited participants, we excluded 20 (25.6%) nonresponders to the first-hand pain placebo manipulation, seven due to technical difficulties with the pain stimulator, five due to inconsistent ratings (e.g., equally high ratings for painful and non-painful stimulation in both placebo and control conditions) and/or extensive movement and one participants because of a spontaneously found brain abnormality.

Our final sample included 22 males and 23 females between 19 and 32 years *(M* ± *SDage* = 23.84 ± 2.73, all strongly right-handed with laterality quotients (LQs) ≥ 80 and normal or corrected-to-normal vision). For the purpose of our study, we only recruited strongly righthanded participants and always induced the placebo in the right hand of all participants to avoid laterality-related problems in our fMRI analyses, increase sample homogeneity and retain comparability of the induction procedure. All participants gave written consent at each outset of their two sessions. Each participant received 50 € for taking part in both sessions and an amount aliquot to their time invested if they dropped out earlier. The study was approved by the ethics committee of the Medical University of Vienna (EK-Nr. 661/2011) and performed in line with the latest revision of the Declaration of Helsinki (2013).

### 2.4 Procedure

Participants came to the MRI scanner, where the experimenter explained that the goal of the study was to investigate brain activity associated with a local anesthetic in the form of a medical gel. First, we performed an individual psychophysical pain calibration. The goal of the calibration was (a) to determine the maximum level of painful, but tolerable stimulation intensity and (b) to specify average subjective intensities on a visual analogue scale (VAS) from 0 = not painful to 8 = extremely painful for very painful (rating of 7), medium painful (4) and not painful, but perceivable (1) stimulation. Participants were asked to rate each stimulation as intuitively but also as accurately as possible. We calibrated the left and right hand individually for each participant (alternating the hand being calibrated first across participants), because previous studies found pain tolerances to vary depending on the hand laterality and dominance (Murray & Safferstone, 1970; Pud et al., 2009). Using the procedure employed by Rütgen, Seidel, Silani, et al. (2015), one electrode was attached to each dorsum of participants’ left and right hands with medical tape. Electrical stimulation (stimulus duration = 500 ms) was administered using the Digitimer DS5 Isolated Bipolar Constant Current Stimulator (Digitimer Ltd, Clinical & Biomedical Research Instruments), one hand at a time, with two rounds going stepwise from very low (0.05 mA) to higher stimulation until the participant indicated the last given stimulus as an ‘extremely painful’ (rating of 8). In a third round, stimuli with seemingly random intensity in the before calibrated range were delivered (although we alternated intensities corresponding to average ratings of 1, 4 and 7). A few seconds break between each stimulation and a few minutes break between each round ensured a reliable, independent rating of each stimulation.

The calibration was followed by the placebo analgesia induction. To this end, a medical student posing as the study doctor introduced the gel as a “powerful local anesthetic”, gave information on its effects and possible side effects, and then applied the placebo gel on the dorsum of the right hand. Participants were then told that a “control” gel without any active ingredient would be applied on the left hand. In fact, both the placebo and control gel contained nearly the same ingredients and no active pharmacological components (exact ingredients can be found in section M.3 in the Supplement). After 15 minutes of waiting time “for the medication to take effect”, we evaluated and amplified the effects of the placebo induction by means of a classic conditioning procedure. As in the calibration, participants were told that they would again receive electrical stimulation to the right or left hand via an attached electrode. Participants thought they would receive stimulation they had rated as very painful before on both hands. This was true for the left/control hand (average intensity rated as 7 during calibration), however, for the right/placebo hand, the experimenter secretly lowered the intensity to medium painful (rating of 4) to suggest pain relief. All participants completed a minimum of two and a maximum of four rounds and received oral feedback by the experimenter after each round. This was done to suggest that participant’s ratings on the left/control hand stayed similar to their average ratings during calibration, but the ratings on the right/placebo hand had decreased. A conditioning round was deemed successful, if all stimuli on the right/placebo hand were rated with lower than 6 and all stimuli on the left/control hand higher than 5. After unsuccessful rounds, stimulation intensity was slightly adjusted, i.e., increased for the left/control hand and/or decreased for the right/placebo hand, in order to increase the contrast between the hands. This adjustment was done covertly, to maintain a participant’s belief that they received equally high intensities on either hand. Importantly, we purposefully chose a within-subjects design and therefore did not directly test whether placebo analgesia leads to a general decrease in somatosensory regions, due to the absence of a control group not undergoing the placebo manipulation.

Afterwards, participants were led into the scanner room and, following general adjustments, completed two runs of another task (~45 minutes, reported in Hartmann et al., 2021), and one run of the picture-based empathy for pain task (~22 minutes) in a fixed order. Upon completion of all tasks, the field map and structural image were acquired. The session was concluded with post-experimental questionnaires and took around four hours in total.

### 2.5 Picture-based empathy for pain task

To measure empathy for everyday painful situations, we adapted an established picture-based empathy for pain task from Jackson, Meltzoff, & Decety (2005). Stimuli depicted everyday situations of one person accidently hurting him-/herself. For each situation, four pictures were taken, each picture always depicting both hands of a person taken from a first-person perspective (for the participant), but showing one out of four different conditions: left/control hand pain, right/placebo hand pain, left/control hand no pain, right/placebo hand no pain (see **Fehler! Verweisquelle konnte nicht gefunden werden.**A for example stimuli and section M.4 in the Supplement for specific information regarding the stimuli creation and task presentation).

A non-preregistered, online validation study was conducted prior to the main study to select 15 situations out of 29 initial ones. To this end, we had recruited an independent sample of 38 right-handed participants (21 females, *M ± SD_age_* = 30.50 ± 6.27). They were asked to rate 116 stimuli (29 situations x 2 intensities x 2 target hands) regarding other-related pain and one’s own unpleasantness when viewing the picture, both on 9-point visual analogue scales from 0 = “not at all” to 8 = “extremely painful/unpleasant”. From the validation study, 15 out of the 29 situations rated as most painful were selected for the main task and evaluated on the following criteria: i) the stimuli for left and right hand did not differ in the measured dimensions and ii) the stimuli in the pain and no pain conditions did differ from each other. To ensure this, we performed two repeated-measures ANOVAs, for pain and unpleasantness ratings, each including the factors *target hand* (left vs. right hand) and *intensity* (pain vs. no pain) and the ratings from the 15 selected situations (see section M.4 and R.1 in the Supplement for description and results regarding the additionally assessed ratings of arousal, valence and realism).

In the main task, 15 situations per condition were shown, resulting in 60 pictures/trials. The 60 images were included in one out of four pseudorandom trial orders previously created (see **Fehler! Verweisquelle konnte nicht gefunden werden.**B for an overview of the trial structure in the task). Each trial consisted of the picture shown centered on a black background for 3500 ms, a jittered waiting period with a white fixation dot on black background for 5000 ± 2000 ms, two rating questions of 4000 ms each played in random order and a jittered inter-trial-interval of 5000 ± 2000 ms, again with a white fixation dot on black background. The hand that the participants had to rate was marked with a black arrow. Tapping into different aspects of empathy (Lamm & Majdandžić, 2015), participants were asked to provide two ratings: (1) “How painful is it for the person in the picture?” and (2) “How unpleasant was it for you to view the picture?”. While the former question aimed to measure pain intensity, i.e., the cognitive-evaluative aspect of the others pain, the latter question aimed to capture the affective-sharing aspect of the empathic experience. Questions were rated on the same 9-point-scale from 0 = not perceivable to 8 = extremely painful/unpleasant.

### 2.6 Data acquisition

The empathy for pain task was implemented using the Cogent 2000 Toolbox Version 1.33 (http://www.vislab.ucl.ac.uk/cogent_2000.php) within MATLAB R2017b (Mathworks). MRI data was acquired using a 3 Tesla Siemens Magnetom Skyra MRI-system with a 32-channel head coil (Siemens Medical, Erlangen, Germany). The functional scanning sequence included the following parameters: Echo time (TE) / repetition time (TR) = 34/1200 ms, flip angle = 66°, multi-band acceleration factor = 4, interleaved ascending acquisition, interleaved multi-slice mode, matrix size = 96×96, field of view = 210 mm, voxel size = 2.2×2.2×2.0 mm^3^, 52 axial slices coplanar to the connecting line between anterior and posterior commissure, and slice thickness = 2 mm. Functional volumes were acquired in one run of ~22 minutes (participants completed three runs in total, with short breaks in between; the data reported here were always collected in the last one of them; data from the other two are reported in Hartmann et al., 2021). To acquire structural images, we used a magnetization-prepared rapid gradient-echo sequence (TE/TR = 2.43/2300 ms, flip angle = 8°, ascending acquisition, single shot multi-slice mode, field of view = 240 mm, voxel size = 0.8×0.8×0.8 mm^3^, 208 sagittal slices, slice thickness = 0.8 mm).

### 2.7 Behavioral data analysis

All behavioral data were processed and statistically analyzed in RStudio Version 3.6.1 (R Core Team, 2019; see section M.5 in the Supplement for information regarding analysis and plotting functions). Preregistered *t*-tests were conducted one-sided due to *a priori*, directional hypotheses. Cohen’s *d*’s for behavioral and fMRI analyses were calculated using the effect size calculation spreadsheet (version 4.2) provided by Lakens (2013).

#### 2.7.1 Manipulation checks

We conducted two manipulation checks to evaluate the strength of the first-hand placebo analgesia effect. First, we asked participants three times during the session how effective they believed the medication to be in reducing their own pain on the treated hand: after the gel application = pre-conditioning, after the conditioning = post-conditioning and after all tasks in the post-experimental questionnaire = post-session. For this, we conducted three paired *t*-tests comparing the three time-points with each other. These analyses were preregistered as exploratory, but we expected an increase in belief between pre- and postconditioning by the conditioning procedure as in Rütgen, Seidel, Silani, et al. (2015).

Secondly, we used first-hand pain ratings from the two runs with the cue-based pain task preceding the picture-based task in the same imaging session. There, participants had received and rated electrical stimulation on either hand. As this measure (which was also one of the four preregistered criteria to identify non-responders) was a reliable indicator of placebo analgesia, we also used it here as an additional manipulation check (see also section M.2 in the Supplement). To this end, we conducted a paired *t*-test using the first-hand pain ratings from that task (calculated as an index of pain - no pain). We further visually inspected the time course of the ratings related to the placebo hand to (a) pinpoint possible decreases of the placebo effect over the course of that task and (b) confirm that the placebo effect was intact and robust right before participants engaged in the picture-based task.

#### 2.7.2 Preregistered analyses

As those manipulation checks, and thus the induction of a first-hand placebo analgesia effect, were deemed successful, we conducted our main analyses to test whether the firsthand effect of placebo analgesia transferred to picture-based empathy for pain. In general, we employed a within-subjects, full-factorial design with two factors of two levels each *(target hand:* left/control vs. right/placebo hand, *intensity:* pain vs. no pain). Two parametric two-factorial (2×2) repeated measures analyses of variance (ANOVAs) were used to analyze the results, where the dependent variables were either the pain or the unpleasantness ratings related to the viewing of the pictures. For each of the ANOVAs, we then compared the two hands on the index scores (pain - no pain), separately for pain and unpleasantness ratings, using paired *t*-tests (run one-tailed for rating_right hand_ < rating_left hand_).

#### 2.7.3 Post hoc analyses

As the pain rating data was not normally distributed (indicated by a significant Shapiro-Wilk normality test), we additionally calculated a Wilcoxon rank-sign test which mirrored the results of our parametric test (see section R.2 in the Supplement). Because of the (absence of predicted) behavioral results, we added two Bayesian paired-samples *t*-tests to evaluate evidence of absence of effects (Keysers et al., 2020). These analyses mirrored the preregistered behavioral analyses, using pain and unpleasantness ratings separately as dependent variables and the default Cauchy (0, .707) prior of .707 as the effect size (this indicates a 50% chance to observe an effect size between −.707 and .707; see e.g., Rouder et al., 2009). In general, a Bayesian *t*-test generates one Bayes Factor that compares relative evidence for the alternative vs. null hypothesis (BF_10_, H_1_ vs. H0; BF_10_ ≤ 3: weak evidence, BF_10_ > 3: moderate evidence; BF_10_ > 30: strong evidence for H_1_) or the opposite, evidence for the null vs. the alternative hypothesis (BF_01_ = 1/BF_10_; BF_01_ ≤ 3: weak evidence, BF_01_ > 3: moderate evidence; BF_01_ > 30: strong evidence for H_0_) (Giolla & Ly, 2019; van Doorn et al., 2020; Wagenmakers et al., 2011). Bayesian analyses were run one-tailed for ratin_gright hand_ < ratin_gleft hand_ in JASP version 0.11.1 (JASP Team, 2019).

Our present behavioral results showed no evidence for a placebo-related decrease of subjective ratings related to the right/placebo hand. However, in another, previously published empathy task employing electrical stimulation, we did find evidence for a “generalized placebo effect”, i.e., a general down-regulation of empathy for pain ratings in both control and placebo hand, independent of the localized placebo induction on only one hand (see also Supplement B of Hartmann et al., 2021). In other words, the placebo manipulation had a general effect on the rating of our stimuli, independent of the target hand. Therefore, we wanted to investigate the existence of such a generalized downregulation by comparing behavioral ratings of the pictures used in our main study to the same pictures used in the validation study, where no placebo manipulation was employed. This was done by calculating two between-subjects ANOVAs, including either the pain or unpleasantness ratings in regard to the pictures, as well as the factors *target hand* (left/control vs. right/placebo hand), *intensity* (pain vs. no pain) and *study* (validation vs. main study). Effects indicating between-study differences were followed up using Welch’s *t*-tests and evaluated two-sided. Note that none of these analyses were preregistered as they resulted from pattern of results and additional insights and results originating between preregistration and data analysis, and should thus be considered as exploratory.

### 2.8 fMRI data preprocessing and analysis

#### 2.8.1 Preprocessing and first-level analysis

Statistical Parametric Mapping (SPM12, Wellcome Trust Centre for Neuroimaging, https://www.fil.ion.ucl.ac.uk/spm/software/spm12/) running on MATLAB Version R2017b (Mathworks) was used for preprocessing and statistically analyzing the MRI data. Preprocessing involved slice timing (reference = middle slice; Sladky et al., 2011), realignment with each participant’s individual fieldmap, coregistration of structural and functional images, segmentation into gray matter, white matter (WM) and cerebrospinal fluid (CSF), spatial normalization, and spatial smoothing (8 mm full-width at half-maximum Gaussian kernel). The first-level design matrix of each participant contained four condition regressors and one for all ratings. For each condition, the onset and viewing duration (3.5 s) of the pictures were modeled as blocks and convolved with SPM12’s standard canonical hemodynamic response function. Six realignment parameters and two regressors modeling WM and CSF were included as nuisance regressors (WM and CSF values were extracted using the REX toolbox by Duff, Cunnington, & Egan, 2007).

#### 2.8.2 Preregistered analyses

To test our hypothesis of a transfer of the first-hand localized placebo effect to empathy for everyday painful situations, we extracted and analyzed brain activation in three regions of interest (ROIs), determined independently from our data based on a meta-analysis on pain empathy (Lamm et al., 2011) and also used in our previous studies (Hartmann et al., 2021; Rütgen, Seidel, Silani, et al., 2015): left AI (x = −40, y = 22, z = 0), right AI (39, 23, −4) and aMCC (−2, 23, 40). Additionally, we analyzed four ROIs in bilateral S1 (left S1: −39, −30, 51; right S1: 36, −36, 48) and S2 (left S2: −39, −15, 18; right S2: 39, −15, 18), taken from independent findings investigating somatosensory pain perception (Bingel et al., 2004, 2007). We created 10 mm spheres around each of the seven coordinates with MarsBaR (Brett et al., 2002) and extracted parameter estimates for each ROI using the first-level contrast images of each participant and condition with REX (Duff et al., 2007). ROI analyses were run in RStudio Version 3.6.1 (R Core Team, 2019). As in the behavioral analysis, we employed the same within-subjects, full-factorial design including two factors (*target hand*, *intensity)* of two levels each, and the additional factor *ROI* with seven levels (lAI, rAI, aMCC, lS1, rS1, lS2, rS2). We first calculated an ANOVA pooling the activation of all seven ROIs and then calculated separate ANOVAs and planned comparisons for each ROI. Each ANOVA was followed up by a planned comparison between the two hands using an index of the activation (pain -no pain). Since Mauchly’s test for sphericity was statistically significant in the pooled ANOVA for the main effect of ROI and all interactions with the factor ROI, we reported the results of this ANOVA using Greenhouse Geisser sphericity correction. After observing a significant main effect of and interactions with the factor ROI in the initial threeway ANOVA, we went on with our preregistered plan and computed separate ANOVAs and planned comparisons for each ROI (one-sided tests for activity_right hand_ < activity_control hand_). To control for multiple comparisons, we corrected the calculated *t*-tests using Bonferroni correction by dividing the *α* error probability by the number of ROIs (for tests in affective ROIs: *p* = .05 / 3 ROIs = .017; somatosensory ROIs: *p* = .05 / 4 ROIs = .013).

#### 2.8.3 Post hoc analyses

Again, due to the (absence of predicted and exploratory evidence for opposite) results, we ran Bayesian paired-samples *t*-tests for each of the seven ROIs in JASP. The analyses of left and right S1 and S2 mirrored the preregistered fMRI analyses, and were run one-sided for activity_right hand_ < activity_left hand_ using the default Cauchy (0, .707) prior of .707 as the effect size. However, the analyses for the three affective ROIs (lAI, rAI and aMCC) were run one-sided for activity_right hand_ > activity_left hand_ to investigate evidence for the opposite effect.

## 3 Results

### 3.1 Behavioral results

#### 3.1.1 Validation study

The aim of the validation study was to ensure that i) the stimuli for left and right hand did not differ and that ii) the stimuli in the pain and no pain conditions did differ from each other. We calculated five repeated measures ANOVAs for each of the five rating scales (pain, unpleasantness, realism, arousal and valence) including the factors *target hand* (left vs. right hand) and *intensity* (pain vs. no pain). The results showed significant main effects of intensity in all ANOVAs (see Table S1 in the Supplement). Participants judged painful stimuli as significantly more painful (*F*(1,37) = 538.24, *p* < .001, *η*^2^ = 0.89; Figure 2A) and unpleasant (*F*(1,37) = 253.65, *p* < .001, *η^2^* = 0.75; Figure 2B) as their non-painful counterparts. These results successfully demonstrated the validity of our created task stimuli.

**Figure 1.**
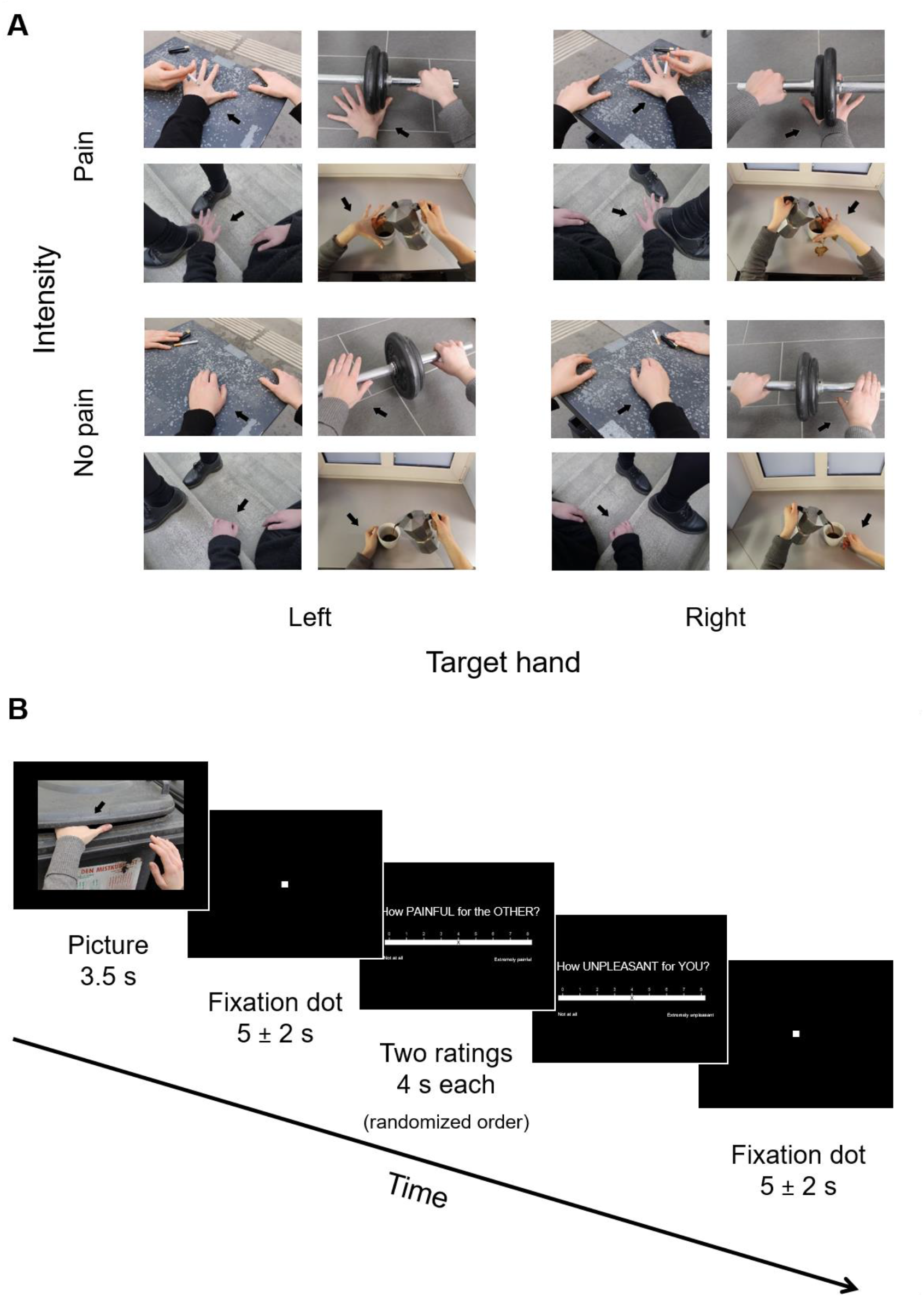
A) Task stimuli showed everyday situations where one person (ostensibly accidentally) hurt him-/herself and varied depending on the target hand (left vs. right) and intensity (pain vs. no pain). Both hands were shown in all stimuli, but black arrows expressly indicated the hand to attend to and to rate in the trial. B) Overview of the picture-based empathy for pain task. In a 2×2 within-subjects design, pictures depicted either painful or non-painful everyday situations, and participants were asked to focus on either the left or right hand (i.e., the one corresponding to their own control or placebo hand, respectively). The hand to be attended to was marked again with a black arrow.

**Figure 2.**
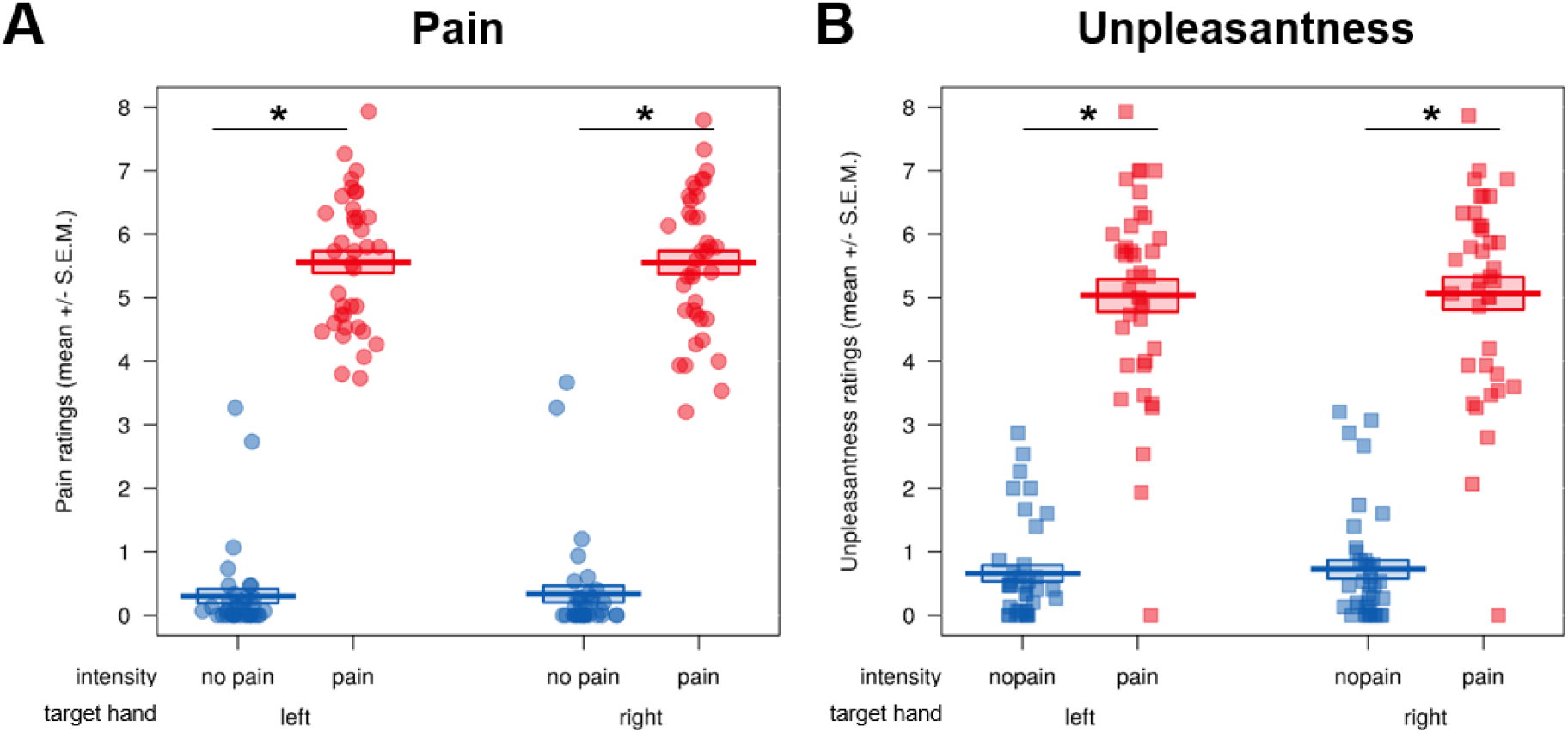
Behavioral results of the validation study. Participants rated picture stimuli of everyday painful and non-painful situations, separately for the left and right hand, and according to the following dimensions, all on 9-point Likert scales: A) pain (0 = “not at all” to 8 = “extremely painful”), B) unpleasantness (0 = “not at all” to 8 = “extremely unpleasant”). In both 2×2 ANOVAs using the factors target hand (left vs. right) and intensity (pain vs. no pain), we observed significant main effects of intensity, demonstrating that painful stimuli were rated as significantly more painful and more unpleasant than their non-painful counterparts. We found no significant differences in our variables between the two hands (main effect of target hand) or any interaction between intensity and target hand. Ratings for arousal, valence and realism can be found in Figure S1 in the Supplement. The validation study was not preregistered, but conducted before (the preregistration of) the main study.

#### 3.1.2 Manipulation checks

Next, we conducted two manipulation checks to evaluate the existence of a first-hand placebo analgesia effect. In the first check measuring beliefs in the effectiveness of the “medication” over the course of the session, means ± standard errors of the mean *(SEM)* were 6.64 ± 0.27 pre-conditioning, 8.06 ± 0.19 post-conditioning and 6.71 ± 0.39 post-session. The increase in effectiveness beliefs as a result of the conditioning procedure was significant (pre- vs. post-conditioning: *t*(44) = 5.91, *p* < .001, *M_diff_* = 1.42, 95% CI_meandiff_ [0.94, 1.91], see Figure 3A).

**Figure 3.**
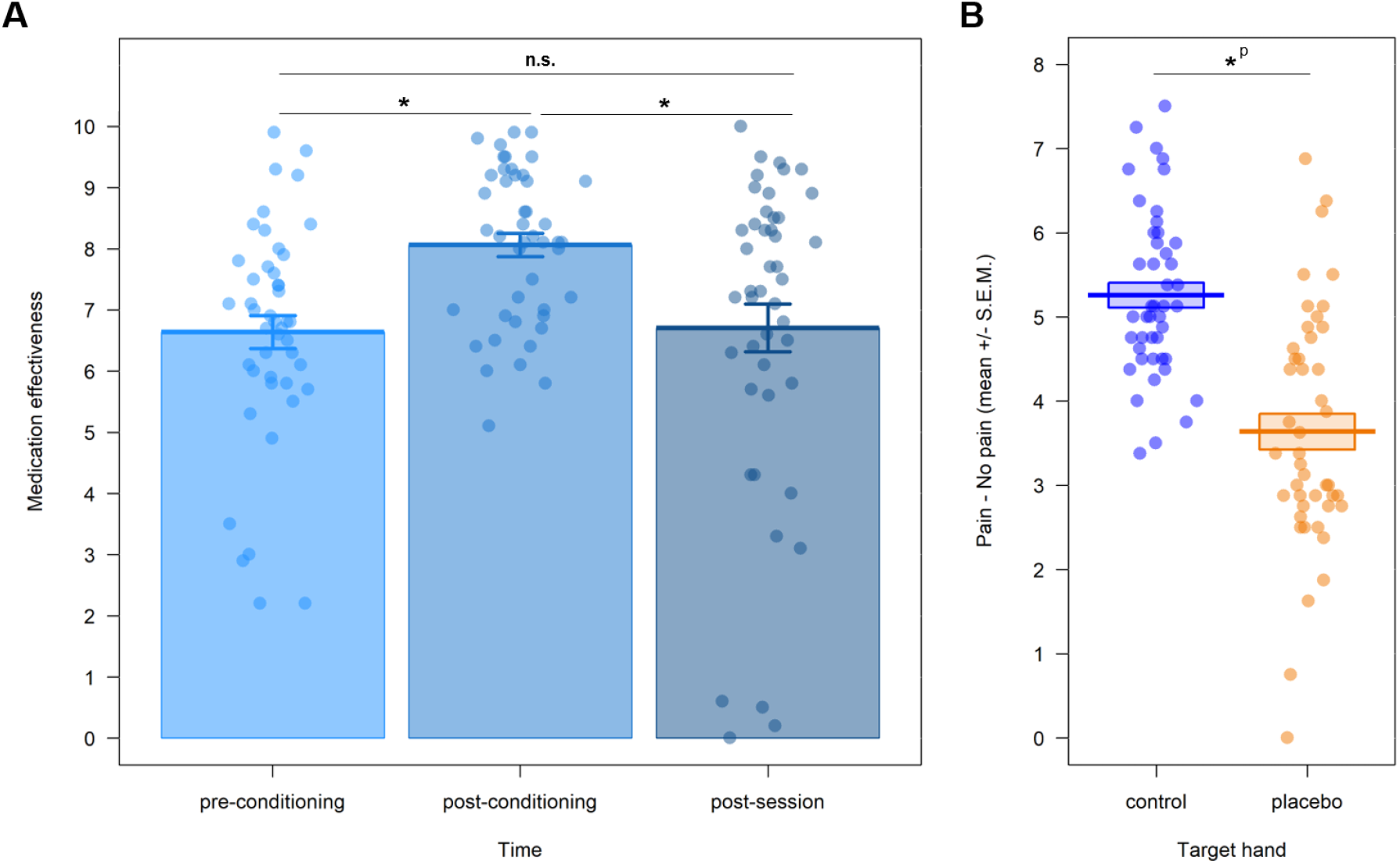
Behavioral manipulation checks to evaluate the strength of the first-hand placebo effect. Displayed are means and standard errors of the mean. A) We evaluated beliefs in the effectiveness of the administered gel at three times during the experiment - directly after the application by the medical cover (pre-conditioning), after the conditioning procedure (post-conditioning) and during the debriefing at the end of the session (post-session). This revealed a significant increase of beliefs after the conditioning. Although the beliefs significantly decreased until the end of the session, they did not drop significantly lower than initial belief levels, which were already high to begin with. B) We further evaluated first-hand pain ratings from another task done in the same session where individually calibrated, painful and non-painful electrical stimulation was delivered to the right and left dorsum of participant’s hands (for a thorough description of this task, see Hartmann et al., 2021); displayed here as an index of the ratings for painful – non-painful stimulation). Here we observed a significant placebo analgesia effect. * *p* < .05; S.E.M. = standard error of the mean; *p* = preregistered. Images taken from Figure 4 and Figure A1 in Hartmann et al. (2021).

Participants’ beliefs dropped after the completion of the task (post-conditioning vs. post-session: *t*(44) = −3.80, *p* < .001, *M_diff_* = −1.36, 95% CI_meandiff_ [−2.07, −0.64]), but did not drop lower than the initial effectiveness belief after gel application after the whole session (preconditioning vs. post-session: *t*(44) = 0.16, *p* = 0.875, *M_diff_* = 0.07, 95% CI_meandiff_ [−0.78, 0.92]).

In the second check, we found a significant difference between the participant’s placebo-treated vs. control-treated hands for first-hand pain ratings in the task preceding the picture-based task, with a very high effect size (*t*(44) = 9.49, *p* < .001 one-sided, *M_diff_* = 1.619, 95% CI_meandiff_ [1.28, 1.96], Cohen’s *d_z_* = 1.42; see Figure 3B). This difference was related to significantly lower pain ratings for the right hand (*M* ± *SEM* = 3.64 ± 0.21), i.e., the hand in which placebo analgesia was induced, compared to the left, control-treated hand (*M* ± *SEM* = 5.26 ± 0.15). Importantly, visual inspection of those first-hand pain ratings showed no sign of decrease over time, suggesting a stable and robust placebo effect right before participants engaged in the picture-based task reported here.

In sum, the two manipulation checks showed a) a strong belief in the effectiveness of the placebo gel that was highest right before entering the scanner, and b) lower subjective, first-hand pain ratings in a task that was done right before the picture-based empathy task we focus on here.

#### 3.1.3 Preregistered analyses

Then, we went on to test our hypothesis and evaluate the existence of a transfer of the lateralized first-hand placebo analgesia effect to empathy for everyday painful situations in the main study. To this end, we calculated two repeated-measures ANOVAs (preregistered, see Tables S2 and S3 in the Supplement). The first repeated-measures ANOVA using the pain ratings of the pictures revealed main effects of target hand (*F*(1,44) = 5.42, *p* = .025 two-sided) and intensity (*F*(1,44) = 1348.88, *p* < .001 two-sided), indicating that there were significantly higher pain ratings for the left vs. the right hand *(M_right_* ± *SEM* = 3.02 ± 0.30; *Men* ± *SEM* = 3.13 ± 0.30), and significantly higher pain ratings for painful vs. non-painful pictures *(M_pain_* ± *SEM* = 5.78 ± 0.09; *M_nopain_* ± *SEM* = 0.37 ± 0.07; see Figure 4 for an overview of all ratings). However, we did not observe a target hand x intensity interaction (*p* = .711 twosided), which would have shown evidence for pain-specific effects of the placebo manipulation. Paired comparisons mirrored those results, with no difference between the ratings (calculated as an index of painful - non-painful stimuli) of left (*M* ± *SEM* = 5.38 ± 0.17) and right (*M* ± *SEM* = 5.42 ± 0.14) hands (*t*(44) = −0.37, *p* = .823 one-sided, *M_diff_* = −0.037, 95% CI_meandiff_ [−0.24, 0.16], Cohen’s *d* = 0.03).

**Figure 4.**
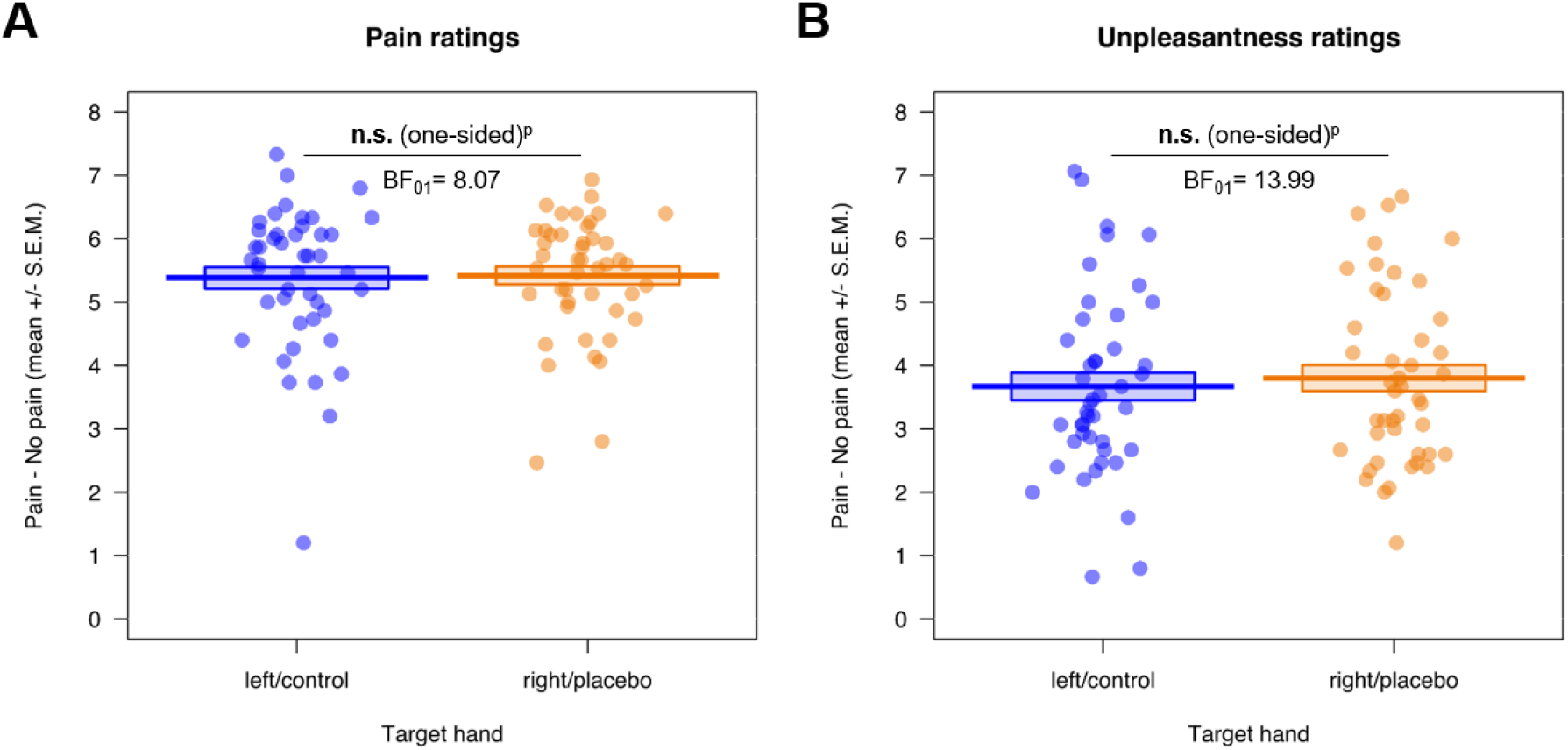
Preregistered behavioral results of the picture-based empathy for pain task, where participants rated pictures of others in painful and non-painful everyday situations (displayed here as an index of the ratings for painful – non-painful control stimuli). We observed no evidence for a transfer of the first-hand placebo effect induced using electrical pain to neither A) empathy for pain nor B) unpleasant-ness ratings, using a one-sided test of right/placebo hand < left/control hand. n.s. = not significant, S.E.M. = standard error of the mean; BF_01_ = evidence for the null compared to the alternative hypothesis (also calculated one-sided); ^p^ = preregistered.

Results were similar in the second repeated-measures ANOVA using the unpleasantness ratings of the pictures. Here, we again observed main effects of target hand (*F*(1,44) = 15.56, *p* < .001 two-sided) and intensity (*F*(1,44) = 327.89, *p* < .001 two-sided), but opposite to the effect found in the pain ratings, there were significantly higher unpleasantness ratings for the right vs. the left hand *(M_right_* ± *SEM* = 2.29 ± 0.23; *M_left_* ± *SEM* = 2.14 ± 0.23), and significantly higher unpleasantness ratings for painful vs. non-painful pictures *(M_pain_* ± *SEM* = 4.08 ± 0.15; *M_nopain_* ± *SEM* = 0.35 ± 0.07). Again, we observed no target hand x intensity interaction (*p* = .160 two-sided), which would again have shown evidence for pain-specific effects. Paired comparisons (using an index of the ratings for painful – non-painful stimuli) showed no difference between the two hands (left: *M* ± *SEM* = 3.67 ± 0.22; right: 3.80 ± 0.21) for unpleasantness ratings (*t*(44) = −1.43, *p* = .960 one-sided, *M_diff_* = −0.13, 95% CI_meandiff_ [−0.32, 0.05], Cohen’s *d* = 0.09).

#### 3.1.4 Post hoc analyses

Complementing the above results, the two post hoc one-sided Bayesian *t*-tests showed moderate to strong evidence for absence of the investigated effect, both for pain (BF01 = 8.07) and unpleasantness ratings (BF01 = 13.99) (see Figure 4) of the main study.

When comparing the main study’s results to the one of the validation study (who had not included a placebo analgesia induction) in order to investigate a generalized effect of this manipulation in the main study, we observed main effects of target hand (*F*(1,81) = 4.00, *p* q .048 two-sided) and intensity (*F*(1,81) = 1649.51, *p* < .001 two-sided), but no main effect or interactions with study (for the full ANOVAs see Tables S4 and S5 in the Supplement). The main effect of target hand showed that pictures of the right hand (*M ± SEM* = 3.04 ± 0.22) were rated as more painful than pictures of the left hand (*M ± SEM* = 2.98 ± 0.22) independent of the intensity or study, and that painful pictures (*M ± SEM* = 5.68 ± 0.07) were rated as more painful than non-painful pictures (*M ± SEM* = 0.35 ± 0.06) independent of the target hand or study.

Looking at the unpleasantness ratings, however, we observed main effects of study (*F*(1,81) = 11.37, *p* = .001 two-sided), target hand (*F*(1,81) = 11.78, *p* < .001 two-sided) and intensity (*F*(1,81) = 576.74, *p* < .001 two-sided). Again, the main effect of target hand showed that pictures of the right hand (*M ± SEM* = 2.57 ± 0.18) were rated as more unpleasant than pictures of the left hand (*M ± SEM* = 2.46 ± 0.18) independent of the intensity or study, and that painful pictures (*M ± SEM* = 4.53 ± 0.12) were rated as more unpleasant than nonpainful pictures (*M ± SEM* = 0.50 ± 0.06) independent of the target hand or study. Interestingly, the main effect of study showed that, in general, pictures were rated as inducing more self-related unpleasant affect in the validation study (*M ± SEM* = 2.87 ± 0.20) compared to the main study (*M ± SEM* = 2.21 ± 0.16). We therefore followed up on the main effect of study by comparing the unpleasantness ratings for the two hands for the two studies, separately for pain and no pain (see Figure 5). Here we observed that, for both intensities, the average unpleasantness rating for either hand in the pictures was significantly higher in the validation compared to the main study (pain: *p* = .004, *M_val_ ± SEM* = 5.05 ± 0.25, *M_main_ ± SEM* = 4.08 ± 0.21; no pain: *p* = .038, *M_val_ ± SEM* = 0.69 ± 0.14, *M_main_ ± SEM* = 0.35 ± 0.09).

**Figure 5.**
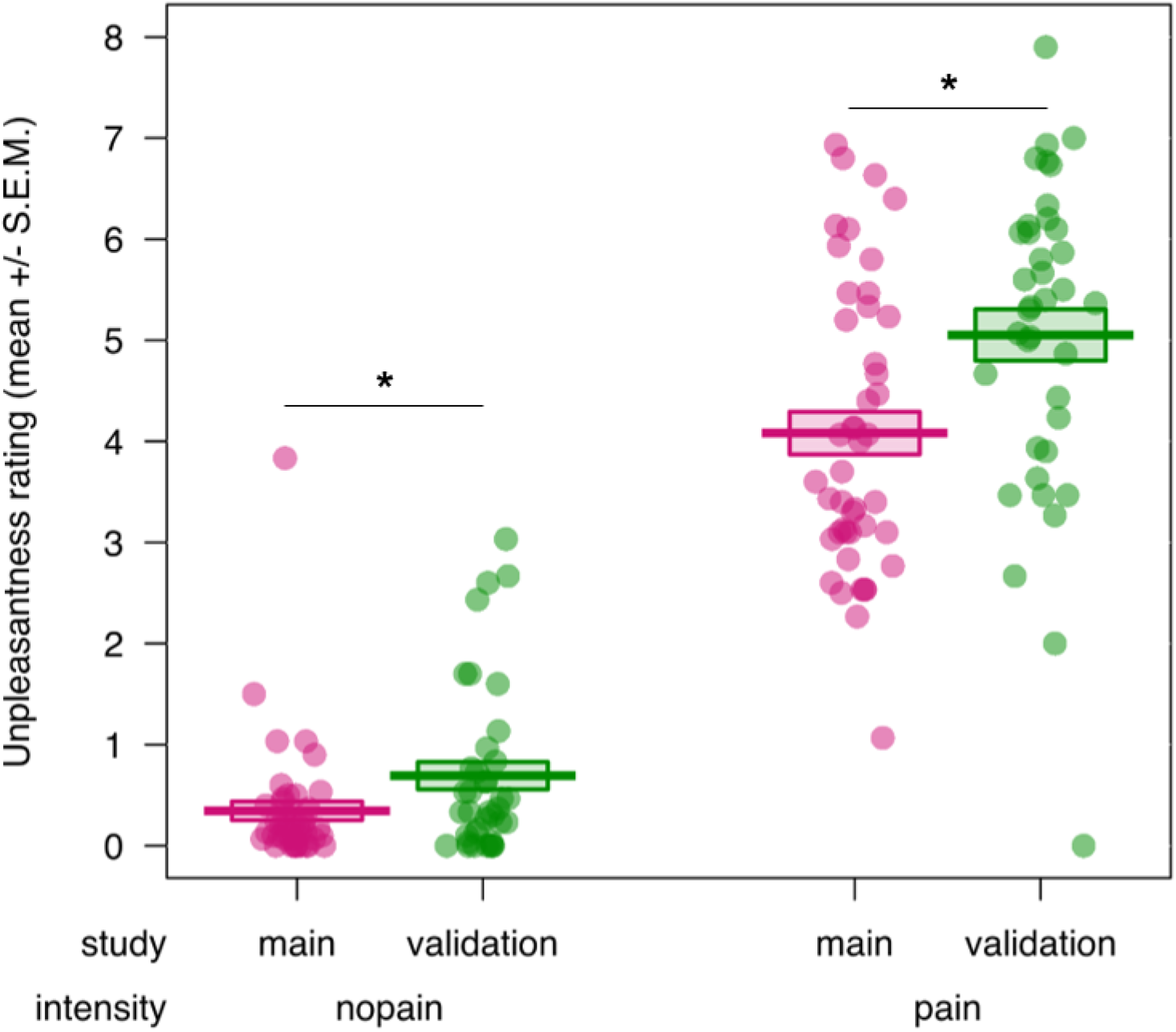
Post hoc comparison of unpleasantness rating to pictures of everyday painful situations between the validation and the main study (displayed here as an average of the ratings for stimuli depicting the right and left hand). We observed evidence for a significant, generalized reduction of unpleasantness in the main compared to the validation study, for pain as well as no pain stimuli. S.E.M. = standard error of the mean.

In sum, the behavioral results indicated no transfer of the placebo analgesia effect induced for first-hand electrical pain to empathy for pain or one’s own unpleasantness during everyday situations. In other words, participants rated the pain intensity of other people in everyday situations as well as their own unpleasant affect when viewing such pictures equally high and independent of the placebo induction on the right hand. These results were confirmed by the post hoc Bayesian analyses. Interestingly, exploratory analyses indicated a possible down-regulatory effect of the placebo induction on unpleasantness ratings to the pictures in general, independent of the rated hand.

### 3.2 fMRI results

#### 3.2.1 Preregistered analyses

After successfully inducing a first-hand placebo analgesia effect in our participants, we went on to test our preregistered main hypothesis for a transfer of this effect to empathy for everyday painful situations using a ROI approach. To this end, we extracted parameter estimates in three affective (bilateral AI, aMCC) and four somatosensory ROIs (bilateral S1 and S2). However, using our preregistered one-sided tests and hypothesizing a reduction of brain activation for pictures corresponding to the placebo hand, we did not find results confirming our hypotheses (see Table 1 for an overview of all planned comparisons).

**Table 1.**
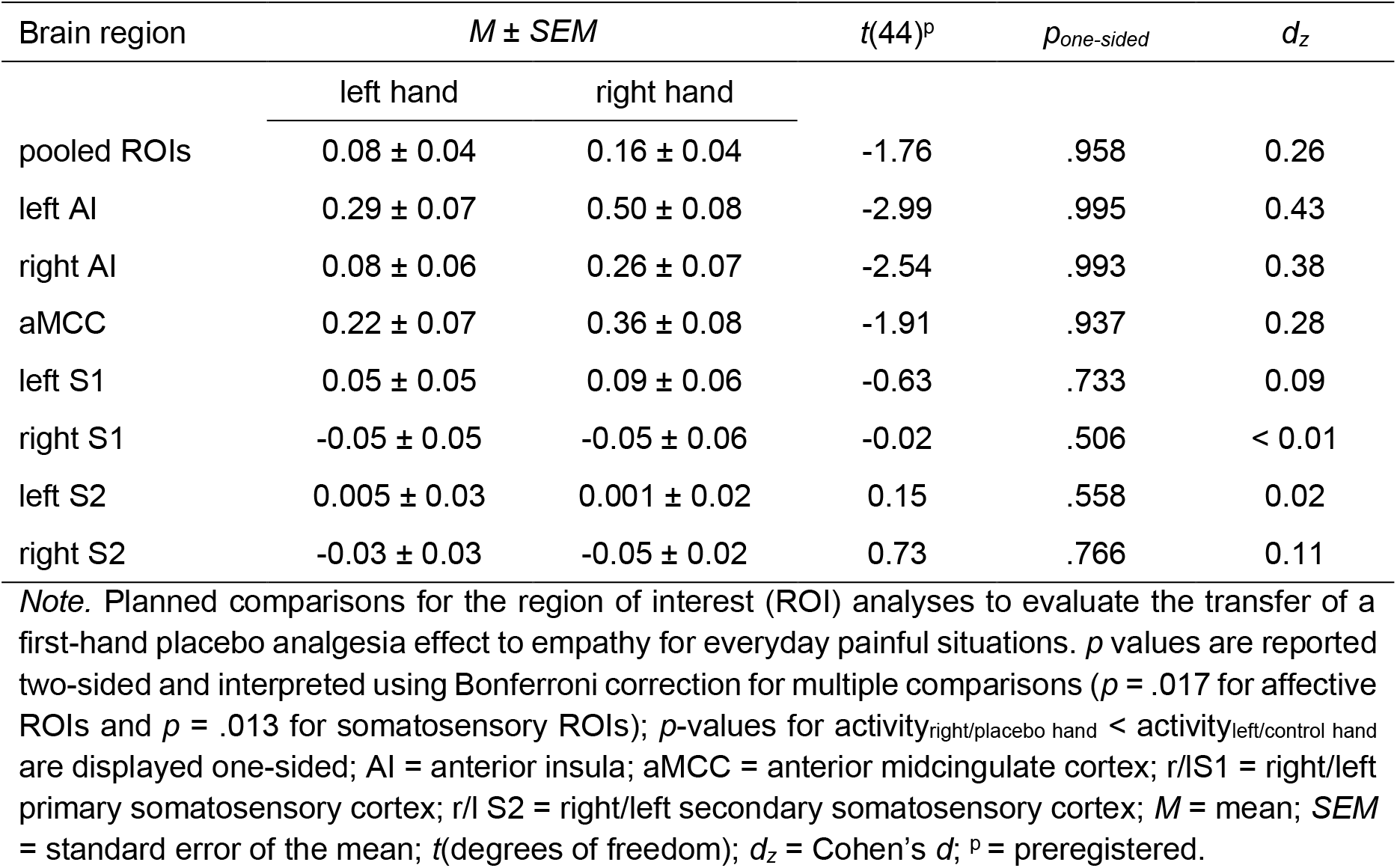
Main ROI analyses testing for placebo effects via paired t-tests.

The pooled ANOVA showed significant main effects of intensity and ROI, a trend for hand x intensity (*p* = .086 two-sided) as well as all interactions involving the factor ROI (see Table S6 in the Supplement for the full ANOVA table). The planned comparison encompassing activation of all ROIs revealed a non-significant result (*p* = .958 one-sided, *M_diff_* = −0.08, 95% CI_meandiff_ [−0.17, 0.01]), with no difference in activation for the placebo and the control hand. As preregistered, and due to a significant main effect of ROI as well as significant interactions with the factor ROI, we went on to calculate single ANOVAs and complementary t-tests for each ROI separately.

Regarding the affective ROIs, we observed no evidence for lower placebo-induced hemodynamic activity in regard to pictures displaying the right/placebo compared to the left/control hand (left AI: *p* = .995 one-sided, *M_diff_* = −0.22, 95% CI_meandiff_ [−0.37, −0.07]; right AI: *p* = .993 one-sided, *M_diff_* = −0.18, 95% CI_meandiff_ [−0.32, −0.04]; aMCC: *p* = .937 one-sided, *M_diff_* = −0.14, 95% CI_meandiff_ [−0.29, 0.01] also see Figure 6 here and Tables S7-S9 in the Supplement).

**Figure 6.**
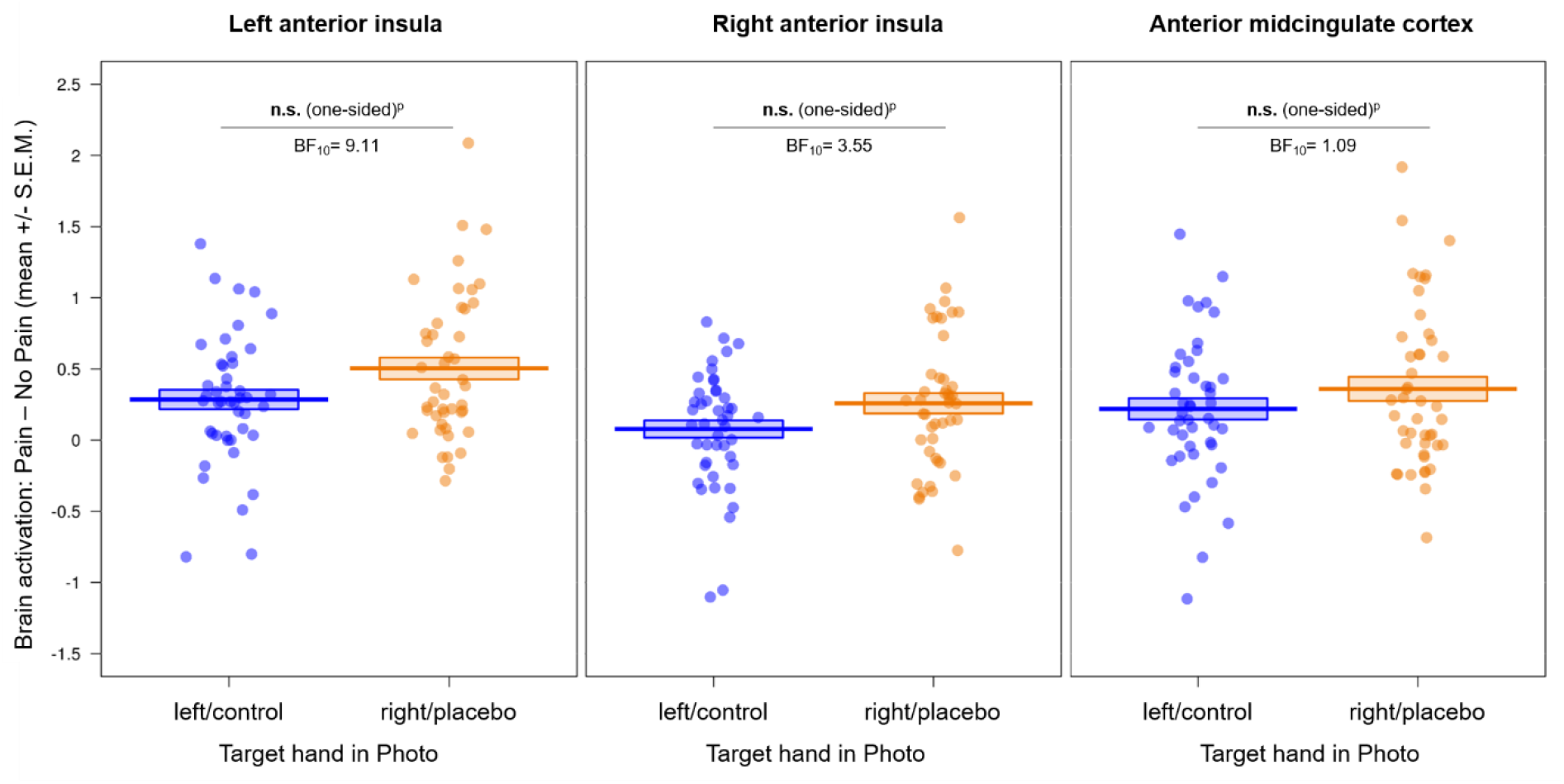
Affective region of interest (ROI) results of the empathy for pain task (displayed here as an index of activity related to painful – non-painful control stimuli). We observed no evidence for a reduction of brain activity in affective regions related to empathy for pain by the placebo manipulation, using a one-sided test of right/placebo hand < left/control hand. Data inspection revealed the opposite effect, i.e., higher activity in affective brain regions during empathy for pain in the placebo compared to the control condition. n.s. = not significant, S.E.M. = standard error of the mean; BF_10_ = evidence for the alternative compared to the null hypothesis; ^p^ = preregistered.

In the somatosensory ROIs, we again found no evidence for our preregistered hypotheses, as there were no differences in brain activation for the two target hands, neither in S1 nor S2 (see Figure 7 and Tables S10-S13 in the Supplement). More specifically, we did not observe the hypothesized lower brain activity related to right hand compared to left hand pictures (left S1: *p* = .733 one-sided, *M_diff_* = −0.05, 95% CI_meandiff_ [−0.19, 0.10]; right S1: *p* = .506 one-sided, *M_diff_* < −0.001, 95% CI_meandiff_ [−0.13, 0.13]; left S2: *p* = .558 one-sided, *M_diff_* = 0.004, 95% CI_meandiff_ [−0.05, 0.06]; right S2: *p* = .766 one-sided, *M_diff_* = 0.02, 95% CI_meandiff_ [0.04, 0.08]).

**Figure 7.**
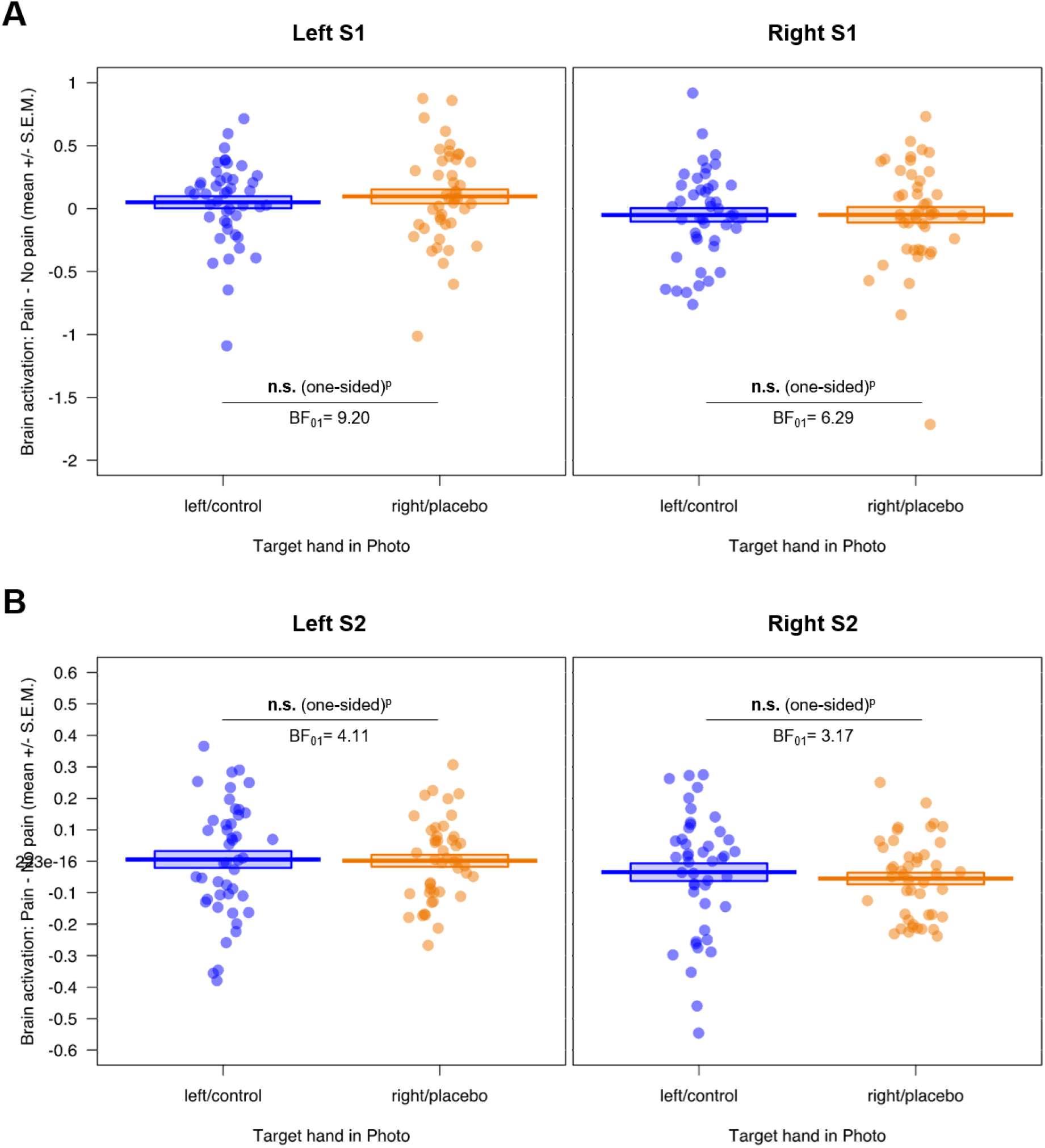
Somatosensory region of interest (ROI) results of the empathy for pain task for A) bilateral S1 and B) bilateral S2 (displayed here as an index of activity related to painful – non-painful control stimuli). We observed no evidence for a reduction of brain activity in somatosensory regions related to empathy for pain by the placebo manipulation, using a one-sided test of right/placebo hand < left/control hand. n.s. = not significant, S.E.M. = standard error of the mean; S1/S2 = primary/secondary somatosensory cortex; BF_01_ = evidence for the null compared to the alternative hypothesis; ^p^ = preregistered.

#### 3.2.2 Post hoc analyses

Inspection of the data revealed an opposite pattern in bilateral AI and aMCC, whereby activation seemed to be higher for right hand compared to left hand pictures. Post hoc Bayesian analyses of the seven ROIs can be found in Table S14 in the Supplement. Importantly, those only showed moderate evidence for H_1_ compared to H_0_ for lAI (BF_10_ = 9.11) and rAI (BF_10_ = 3.55), i.e., higher hemodynamic activity for right vs. left hand pictures, corresponding to participant’s placebo hand, but weak evidence for H1 in aMCC (BF_10_ = 1.09). In contrast, the Bayesian factors (BF_01_) for the somatosensory ROIs showed moderate evidence for the absence of a difference in activity.

In sum, the fMRI results, similar to the behavioral results, did not confirm our predictions of first-hand placebo analgesia reducing empathy for pain in both affective and somatosensory brain regions. However, exploratory post-hoc analyses suggested an opposite effect, i.e., higher hemodynamic activity related to right hand pictures in bilateral AI.

## 4 Discussion

This preregistered study aimed to investigate the role of sensory-discriminative brain responses in empathy for everyday painful situations. To this end, we induced a localized placebo analgesia effect in the right hand of 45 participants using a placebo gel (with the left hand acting as a control) and then measured brain activity with fMRI during an empathy for pain task focusing on visual depictions of everyday painful situations occurring to left or right hands.

Contrary to our preregistered predictions, we did not observe a localized reduction of self-reported empathy for pain or unpleasantness ratings related to the hand laterality as a result of the first-hand placebo analgesia induction. The neuroimaging results mirrored the behavioral findings, as we did not observe evidence for lower hemodynamic activity in any of our affective or somatosensory ROIs as a result of the placebo. These null results were bolstered by three additional findings: First, pain and unpleasantness ratings in the validation study showed a reliable differentiation of painful vs. non-painful stimuli, but no differences on those dimensions for pictures displaying the left vs. right hand, speaking for the validity and comparability of our stimuli. Second, two initial manipulation checks showed an increase in the belief about medication effectiveness through the conditioning procedure, as well as a significant difference between first-hand pain ratings of the right and left hand in another task done in the same session (Hartmann et al., 2021). Together, these results demonstrated the successful induction of a localized, first-hand placebo analgesia effect, thus laying a strong foundation to investigate their possible transfer to the picture-based empathy task. Third, Bayesian analyses using the behavioral and brain data showed moderate to strong evidence for the absence of a transfer of lateralized placebo analgesia for first-hand pain to empathy for pain in our picture-based task.

Our results shed a different light on previous studies reporting an overlap of first-hand and empathy for pain in the sensory-discriminative component (Lamm et al., 2011 for a meta-analysis) and highlighting the role of somatosensory representations in empathy for pain (Avenanti et al., 2005; Bufalari et al., 2007; Riečanský & Lamm, 2019). However, it is important to note that unlike before, our study specifically targeted somatotopic matching of self- and other-related pain states by using a localized placebo manipulation targeted only at the right hand. One possible explanation for the here reported absence of somatosensory sharing could be that the general processing of another’s pain might have a higher weight than processing the exact location and source of that pain, leading, in turn, to less involvement of the somatosensory network. Indeed, we cannot exclude a general downregulation of somatosensory brain regions by placebo analgesia in the present study, as we did not have a between-subjects control group who was not under the influence of the placebo. Our findings further relate to studies investigating individuals with congenital insensitivity to pain (CIP), a patient group which is impaired in their first-hand and thus somatosensory pain processing (Danziger et al., 2006, 2009). CIP patients still show intact empathic responses despite this constraint, including the aMCC and AI engagement seen in neurotypical controls, as well as ventromedial prefrontal and posterior cingulate cortices. Interestingly, those results were found using mainly picture-based measurements of empathy for pain (verbally presented imaginary painful situations, images of body parts and facial expressions and video material of painful situations), as in the present study. Indeed, previous research from our lab found evidence for affective sharing only in a cue-based empathy task employing electrical stimulation in three consecutive studies (Rütgen et al., 2018; Rütgen, Seidel, Riečanský, et al., 2015; Rütgen, Seidel, Silani, et al., 2015). Furthermore, our results are in line with the findings from the cue-based task performed as part of the present study, before the picture-based task reported here. There, we also did not observe evidence for somatosensory sharing employing a task with electrical pain but focusing the attention of participants more specifically to the body part in pain in the same session (Hartmann et al., 2021). The fact that we report no evidence for somatosensory sharing in two separate tasks, but in the same participants, strengthens our conclusion of a less consistent involvement of the sensory-discriminative compared to the affective component in empathy for pain.

We further report exploratory evidence for generally lower unpleasantness as a result of the placebo analgesia induction in the main study, independent of the displayed hands (compared to our prior validation study without such analgesia). Averaging over both hands, we demonstrated significantly higher unpleasantness ratings in the validation study, compared to the main study. This indicates that the placebo induction might indeed have had an effect on the participants, but not in the localized way we expected. Instead, unpleasantness ratings were generally lower in the main compared to the validation study independent of the hand laterality in the stimuli. This finding would be in line with our past work (Rütgen, Seidel, Silani, et al., 2015) showing effects of a generalized placebo analgesia induction on affective brain networks, and with our previous study where we also observed such a general downregulation of empathic responses (Hartmann et al., 2021). More specifically, there we had observed, within the same sample of participants, a comparable reduction of empathy for pain ratings for both hands instead of the expected localized reduction for one hand only (for a thorough discussion see Supplement B of Hartmann et al., 2021). These results underline the conclusion that the placebo analgesia induction in the present project, encompassing the present study and the one published in Hartmann et al. (2021), may have influenced empathic resonance in a generalized, compared to a localized, way. Although this may not be too surprising, as the same participants underwent this manipulation and did both tasks, it is interesting to note that this general effect was observed consistently over both tasks and is in line with other work from our lab reporting evidence for affective shared representations (Rütgen, Seidel, Silani, et al., 2015). However, this reasoning is only preliminary and should be investigated further in future studies, especially because the validation and main study were not exactly the same: Apart from the 15 painful situations used in the validation and main study, the validation study employed a different set of participants, measured an additional 14 situations and also assessed realism, arousal and valence ratings for each picture. Although we chose the 15 most painful situations for the study comparison, the possibility of anchoring effects depending on the shown stimuli should be the subject of future research.

Furthermore, we found an unexpected opposite effect for bilateral AI, whereby hemodynamic activity was higher for pictures of the right hand (corresponding to the participant’s own placebo hand) compared to the left/control hand. It is important to note, though, that these explorations were not part of our preregistered hypotheses and have to be interpreted with caution. Critically, we did not preregister an analysis testing for this unexpected finding of higher brain activity for pictures relating to the right as opposed to the left hand, as we hypothesized the opposite effect and thus tested one-sided. This caveat, coupled with our null findings and the nature of the picture-based task, which did not allow a comparison to the first-hand experience, limit the conclusions we can make in regard to somatosensory sharing of another’s pain. Although we report null findings regarding our preregistered predictions, we briefly discuss our results in the next section and highlight future research directions.

First, we induced the placebo only in the right hand of all our strongly right-handed participants, thus potentially confounding handedness with our placebo manipulation. It could therefore be that the placebo hand, always being the dominant hand, had specific effects on brain activation related to this hand, such as greater salience or attention. Indeed, Atlas & Wager (2014) discuss the possibility of placebo-induced changes in the insula and ACC changes reflecting attention or positive affective shifts. Future studies should therefore carefully test such laterality-related issues and extend this line of research, e.g., by counterbalancing the placebo analgesia induction and/or adding left-handers to their sample.

Second, previous research regarding first-hand as well as empathy for pain has also highlighted the role of stimulus context and appraisal mechanisms (Atlas et al., 2010; Hein & Singer, 2008 for a review; Lamm, Batson, et al., 2007; Lamm, Nusbaum, et al., 2007). For example, it has been shown that perspective changes can modulate evaluative processes and subsequent brain responses related to pain (Jackson et al., 2006), and that pain perception is shaped by multiple context factors such as expectations or beliefs (Jepma et al., 2018; Jepma & Wager, 2013). In our task, for example, the context varied greatly between the pictures and each situation had to be interpreted differently, possibly requiring increased top-down perspective taking. In this line of reasoning, a failure of self-other distinction could have led to ‘empathic over-arousal’ in the form of higher brain activation in AI in the placebo compared to the control condition. However, to avoid confusion of self- and other-related perspectives, but also mixing up of the hand laterality, we purposefully chose to display the pictures from an egocentric perspective and kept a black frame around the images.

Third, we carefully selected placebo responders by means of four specific criteria (Hartmann et al., 2021; Rütgen, Seidel, Silani, et al., 2015), i.e., only included participants showing a first-hand placebo analgesia effect. Wager et al. (2011) have highlighted a possibly altered process of pain evaluation in strong placebo responders, rather than a blockade of noxious input. Indeed, many previous studies not only find a decrease in pain-related brain regions but an additional increase in affective brain regions such as rostral/pregenual ACC or insula for first-hand pain, suggesting placebo-related modulatory mechanisms (Atlas et al., 2010; Bingel et al., 2006; Petrovic et al., 2002, 2005; see Amanzio et al., 2013 and Atlas & Wager, 2014 for meta-analyses). If the pain is evaluated differently to begin with, this could explain our paradoxical increase of brain activity in bilateral AI, corresponding to processing of right-hand pictures. However, this is only speculative and should be thoroughly tested in future studies, especially since the evidence observed here was only moderate and the majority of studies using placebo analgesia usually report a decrease of activity in insular regions.

Two other important points are task order and time effects. Although we took great care in inducing the placebo effect in a way that it was stable for the whole session, the task reported here was conducted after another task, around 45 minutes after the placebo induction was completed. This could have influenced the strength of the placebo effect and led to a decrease or general change over time. However, both of our manipulation checks showed a persistent effect until the end of the session, visible in comparable beliefs in the effectiveness of the gel from the start of the placebo induction to the end of the session, and in significantly lower first-hand pain ratings for the placebo compared to the control hand in another task done right before the here presented empathy task. Furthermore, a previous study points to self-reinforcing expectancy effects in pain (Jepma et al., 2018). Indeed, the first-hand pain ratings in the placebo condition of the cue-based task completed directly before the picture-based task, showed no evidence for a decrease of the placebo effect over time.

As mentioned before, placebo analgesia in first-hand pain may not only have led to decreased brain activity in regions such as insula, ACC and S2, but also to increased activity in prefrontal regions such as dorsolateral, medial and orbitofrontal cortices as well as inferior frontal gyrus (see e.g., Atlas & Wager, 2014 for a meta-analysis). In line with this, neurons in the medial prefrontal areas might be involved in exerting placebo-related, modulatory control over an automatically triggered, affective response to the sight of highly aversive situations such as our stimuli (Lamm, Nusbaum, et al., 2007). However, a complementary whole brain analysis did not reveal any activation differences for placebo vs. control hand in prefrontal areas in our task, when FWE-corrected at cluster-level. Investigating these brain regions in the context of analgesic effects on empathy for pain more specifically should therefore be the focus of future studies.

In conclusion, although our results suggest a successfully induced, localized placebo analgesia effect for first-hand pain, we did not find evidence for a laterality-specific reduction in affective or somatosensory brain areas, when assessing empathy for everyday painful situations. While not showing a transfer of these lateralized effects, the placebo analgesia induction affected participants’ unpleasantness in a more generalized way, which is in line with previous research. Such results are crucial in casting further light of how empathy is processed in the brain and on the extent and type of influence that our first-hand pain experience has on empathic responding.

## 5 Funding

This work was supported by the uni:docs scholarship of the University of Vienna (to H.H.); the Austrian Science Fund (FWF, W1262-B29); and the Vienna Science and Technology Fund (WWTF, VRG13-007). None of the funders had any role in study design, data collection and analysis, interpretation, writing or decision to publish.

## 6 Acknowledgements

We thank Ronald Sladky for valuable input on a final draft of the preregistration. We would also like to thank the two master students Anna Köstler and Fabian Franken who wrote their theses within this project as well as numerous interns and confederates for helping with the data collection.

## 7 CRediT author statement

**Helena Hartmann:** Conceptualization, Data curation, Formal analysis, Funding acquisition, Investigation, Methodology, Software, Visualization, Writing - original draft, Writing - review & editing. **Federica Riva:** Formal analysis, Supervision, Writing - original draft, Writing - review & editing. **Markus Rütgen:** Formal analysis, Methodology, Supervision, Writing - original draft, Writing - review & editing. **Claus Lamm:** Conceptualization, Formal analysis, Funding acquisition, Methodology, Project administration, Resources, Supervision, Writing - original draft, Writing - review & editing (Brand et al., 2015).

## 8 Declarations of Interest

H.H. works as a researcher and psychologist for MyMind GmbH, a company developing a neurofeedback training game for children with Autism Spectrum Disorder, but this work is in no way related to the present research. The remaining authors declare that they have no financial interests or potential conflicts of interest.

## Notes

https://neurovault.org/collections/9244/

