## Supplementary Material for "Placebo analgesia does not reduce empathy for naturalistic depictions of others’ pain in a somatosensory specific way"

### Supplement

This study is part of a bigger project that has in part already been published, with the only difference being the task that was reported (Hartmann et al., 2021). Most of the information given here in M.1, M.2, M.3 and M.5 is therefore paraphrased from this previous paper.

#### Methods

##### M.1 Sample

Participants were recruited via a database of interested study participants, distributing flyers and advertising on social media. Interested participants first filled out an online questionnaire screening for any exclusion criteria (especially regarding MRI safety) and only eligible participants were invited to the study. Exclusion criteria were past/present enrollment in university studies including psychology, pharmaceutics or (human, veterinary, dental, etc.) medicine, any psychiatric or neurological conditions, past/present long term self-injurious behavior, past or present medical conditions regarding the hands that could possibly interfere with current pain sensitivity (e.g. chronic pain, numbness, trauma, injuries or long-term pain therapy), past/present alcohol or drug abuse, the intake of psychopharmacological medication in the last three months (besides oral contraceptives), left-handedness or no handedness preference, and a weakness in distinguishing left from right. The latter was operationalized by asking participants how often they mix up left and right in their daily life, e.g. while making a turn with the car. We excluded participants answering this question with ‘sometimes’, ‘often’ or ‘always’. Handedness was measured using a combination of two established handedness questionnaires (Büsch et al., 2009; Oldfield, 1971). A laterality quotient (LQ; going from -100 = strongly left-handed to +100 = strongly right-handed) was calculated from the ten most selective items of the two questionnaires, and only individuals with LQs ≥ 72 were invited to the take part (Tran, Stieger, & Voracek, 2014). Each participant was tested twice for any contraindications regarding MRI scanning, once via an online screening questionnaire and once in person at the outset of the scanning session. Excluded participants, such as dropouts and placebo analgesia nonresponders (see also next section S.2), were replaced until the preregistered sample size of 45 participants was reached.

##### M.2 Nonresponder identification

Placebo analgesia nonresponders were determined using four exclusion criteria (the first three criteria taken from (Rütgen et al., 2015): (1) We recorded any strong, verbally expressed doubts regarding the study setup during and after the session; (2) We collected belief scores about the effectiveness of the gel to decrease the participant’s own pain at three time-points during the session (after the gel application = pre-conditioning, after the conditioning = post-conditioning, and in the post-experimental questionnaire = post-session, each rating on a continuous visual-analogue-scale from 0-10 cm). Pre- and post-conditioning ratings that were < 6.66 cm in sum (indicating a general low belief) and/or pre- minus post-conditioning ratings that were > 3.33 cm (indicating a decrease in belief after the conditioning) classified a participant as a nonresponder. (3) We counted the number of conditioning trials needed to show an analgesic response, with four or more trials indicating nonresponding. However, none of the participants needed more than three conditioning trials to show a response to the placebo; (4) During the data collection, we preregistered a fourth measure we had previously overlooked, as it had not been possible in the previous study (which mainly informed the pre-registration) due to a between-subjects design. This involved a first-hand pain task, where we applied short-lasting painful and non-painful electrical stimulation delivered to the right and left hands of the participant, and then collected pain ratings (see also Hartmann et al. (2021) for a more thorough report). Using this fourth criterion, we could directly compare the self-related pain ratings of the two hands and excluded participants who showed higher average first-hand pain on the right/placebo compared to the left/control hand. This allowed us to identify responders with increased certainty, and thus maximize the placebo responsiveness of the sample as well as bolster interpretability of our results. During preregistration of this addendum, the so far collected data had not been observed or analyzed yet.

##### M.3 Procedure

In brief, the study consisted of two parts: First, participants were invited to an initial one-hour session to the Faculty of Psychology and were asked to fill out different trait personality questionnaires on a computer for around one hour. For the second session (average interval of 32.86 ± 29.16 (*M* ± *SD*) days), participants were asked to refrain from alcohol, drugs or medication-intake 24 hours before the testing session as well as from intake of food or any other drink besides water one hour before.

For the placebo induction, a combination of verbal suggestions and classical conditioning techniques was employed. First, a medical student posing as the study doctor (either male or female) did a brief medical screening including a (pseudo) drug-test to increase participant’s belief in the cover story that a real medication would be administered. The placebo gel was presented as a “potent, local anesthetic” with pain-reducing effects on the part of the skin where it is applied to for around 2-3 hours and that its maximum effectiveness would thus last the whole scanning time. Importantly, participants were told that the medical gel blocked pain receptors in a pharmacological way and would thus exert no effects on normal touch sensitivity or non-painful stimulation intensity. This was done to ensure a belief decrease due to an absence of possibly expected skin numbing. In addition, the “medication” was described as already being well established since many years, legally approved, and routinely used in e.g. dental procedures and chronic pain patients. Participants were told that possible side effects in very rare cases could be dry skin and slight skin irritation. The placebo gel was always applied first on the dorsum of the right hand and directly after that the control gel on the left hand, which was described as a normal basic skin cream and representing a basis for the participant’s typical pain perception (the word “placebo” was never mentioned to the participants over the course of the experiment). Participants were told that the application of different gels on each hand was routinely done to ensure equal conditions and make them comparable for later analysis. Both the placebo and control gel contained 0.5 g Carbomer, 0.09 g TRIS, 15 g undiluted Isopropanol, 10 g Propylenglykol, 3 g Glycerol and Aqua pur. ad 100 g, the only difference being 10 g (out of 100 g) Isopropanol in the placebo gel and 10 g basic skin cream (‘Ultrasicc’) instead of the Isopropanol in the control gel. This difference led to a clearly recognizable visual and olfactory distinction between the gels, while still keeping them matched with regard to tactile feeling and hydrating properties. Here we adhered to previously used procedures inducing placebo analgesia by means of topical creams and gels (e.g. Benedetti et al., 1999; Bingel et al., 2006; Geuter et al., 2013; Schenk et al., 2014; Tinnermann et al., 2017). After the gel application on both hands, the participant was led outside the control room to (ostensibly) wait for the medication to take effect, and received detailed instructions regarding the task. Then we asked participants back in the control room and told them that a “pain test” would be used to check the effectiveness of the medication. Before the conditioning and application of new electrodes in the same location as during calibration, excess gel was removed, and the hands were disinfected with 70% isopropyl rubbing alcohol. In the first conditioning round three stimulations were administered, in subsequent rounds four.

##### M.4 Empathy for pain task

The pictures were taken with a Canon EOS 1100D Camera (ISO 6400, 12.2 Megapixel) from an egocentric perspective to facilitate “putting oneself in the shoes of the other”, as no mental rotation of the hand position was required of the participants. As in the study by Jackson et al. (2005), all situations depicted events happening in everyday life, such as jamming the hand in a cupboard or burning your hand in the oven and used different types of pain (mechanical, thermal and pressure). Minor clean-up, image improvements and insertion of black arrows were done with Adobe Photoshop CS3. All pictures were rescaled to 700x525 pixels for the task and displayed on a BOLD screen 32 LCD for fMRI (https://www.crsltd.com/tools-for-functional-imaging/mr-safe-displays/boldscreen-32-lcd-for-fmri/nest/boldscreen-32-technical-specification#npm) which participants watched through a mirror mounted on the head coil. The hand participants rated with was counterbalanced between but kept constant within participants over all tasks in order to avoid any effects in the placebo hand being related to hemodynamic activity during rating.

In the validation study, we additionally asked participants to judge the realism (“Can you imagine this happening to you in real life?”) on a 9-point visual analogue scale from 0 = “not at all” to 8 = “extremely realistic”, as well as arousal and valence of each picture (via 9-point Self-Assessment Manikins (SAM) from “calming” to “exciting” (arousal) and from “negative” to “positive” (valence); see e.g. Bynion & Feldner, 2017).

##### M.5 Data analysis and plotting

The following packages (functions) were used for analyses and plotting in RStudio: plyr (ddply), dplyr (arrange), stats (shapiro.test, t.test, cor.test, aggregate), tidyr (gather, spread), reshape2 (dcast), ez (ezANOVA), yarrr (pirateplot) and ggplot2 (ggplot). All Figures were created in Microsoft PowerPoint.

#### Results

##### R.1 Validation study

In Table S1 and Figure S1, we report the ANOVA results of the validation study. Non-painful stimuli were rated as significantly more calming (*F*(1,37) = 130.97, *p* < .001, *η^2^* = 0.53) and positive (*F*(1,37) = 109.92, *p* < .001, *η^2^* = 0.66) than painful stimuli. Although stimuli were judged as realistic in general, painful stimuli were rated as significantly less realistic as non-painful stimuli (*F*(1,37) = 83.77, *p* < .001, *η^2^* = 0.33).

| Table S1  *Behavioral results of the ANOVAs in the validation study.* | | | |
| --- | --- | --- | --- |
| Tests and effects | *F*_(1,37)_ | *p_(two-tailed)_* | *gen. η^2^* |
| **Pain** |  |  |  |
| target hand | 0.12 | .736 | < 0.001 |
| intensity | 538.24 | < .001 | 0.89 |
| target hand x intensity | 0.41 | .524 | 0.001 |
| **Unpleasantness** |  |  |  |
| target hand | 1.21 | .278 | 0.004 |
| intensity | 253.65 | < .001 | 0.75 |
| target hand x intensity | 0.15 | .701 | < 0.001 |
| **Arousal** |  |  |  |
| target hand | 0.54 | .466 | < 0.001 |
| intensity | 130.97 | < .001 | 0.53 |
| target hand x intensity | 1.59 | .215 | 0.001 |
| **Valence** |  |  |  |
| target hand | 2.28 | .139 | < 0.001 |
| intensity | 109.92 | < .001 | 0.66 |
| target hand x intensity | 1.55 | .221 | < 0.001 |
| **Realism** |  |  |  |
| target hand | 1.99 | .166 | < 0.001 |
| intensity | 83.77 | < .001 | 0.33 |
| target hand x intensity | 0.49 | .490 | < 0.001 |

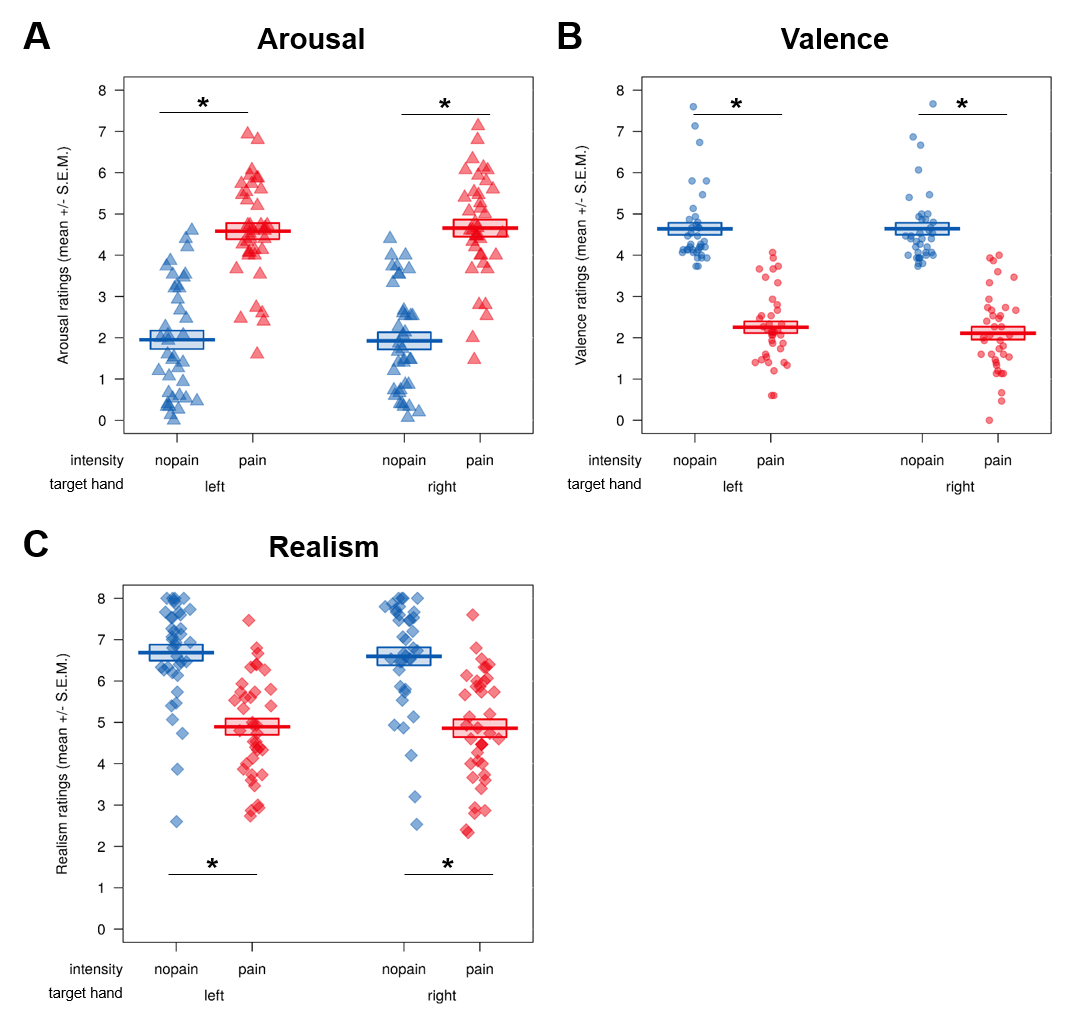

Figure S1. Behavioral results of the validation study. Apart from pain and unpleasantness, participants rated pictures of everyday painful/non-painful situations on three additional dimensions, separately for the left and right hand, all on 9-point Likert scales: A) arousal (0 = “calming” to 8 = “exciting”), B) valence (0 = “negative” to 8 = “positive”) and C) realism (0 = “not at all” to 8 = “extremely realistic”). In all 2x2 ANOVAs using the factors target hand (left vs. right) and intensity (pain vs. no pain), we observed significant main effects of intensity, demonstrating that painful stimuli were rated as significantly less realistic, more arousing and more negative than their non-painful counterparts. We found no significant differences in our variables between the two hands (main effect of target hand) or any interaction between intensity and target hand.

Importantly, although the validation study showed that stimuli were judged as realistic in general, painful stimuli were rated as significantly less realistic than non-painful stimuli. This is no surprise, as we asked how realistic participants imagine that the situations happen to them in everyday life and non-painful situations occur more often. Crucially, we did not find any significant differences in pain, unpleasantness, realism, arousal or valence between pictures focusing on the right or left hand, showing that any possible differences in realism affected the processing of pictures related to either hand in a similar way.

##### R.2 Main behavioral analyses

Because the assumption of normality was violated for the pain rating data in the main study, we additionally calculated a Wilcoxon rank-sign test using the pain ratings. Similar to the preregistered ANOVA and *t*-test above, this did not reveal a statistically significant difference in pain ratings between the two hands (*p* = .293 one-tailed, Cohen’s *d* = 0.03). Below we report the full ANOVA tables of behavioral results of the main study.

| Table S2  Behavioral results of the ANOVA using pain ratings. | | | |
| --- | --- | --- | --- |
| Effects | *F*_(1,44)_ | *p_(two-tailed)_* | *gen. η^2^* |
| hand | 5.42 | .025 | 0.004 |
| intensity | 1348.88 | < .001 | 0.93 |
| hand x intensity | 0.14 | .711 | < 0.001 |

| Table S3  Behavioral results of the ANOVA using unpleasantness ratings. | | | |
| --- | --- | --- | --- |
| Effects | *F*_(1,44)_ | *p_(two-tailed)_* | *gen. η^2^* |
| hand | 15.56 | < .001 | 0.004 |
| intensity | 327.89 | < .001 | 0.74 |
| hand x intensity | 2.04 | .160 | < 0.001 |

##### R.3 Study comparison

Here we report the full ANOVA tables of the exploratory comparison of pain and unpleasantness ratings between the validation and the main study.

| Table S4  Behavioral results of the study comparison ANOVA using pain ratings. | | | |
| --- | --- | --- | --- |
| Effects | *F*_(1,81)_ | *p_(two-tailed)_* | *gen. η^2^* |
| study | 1.19 | .277 | 0.01 |
| hand | 4.00 | .049 | 0.001 |
| intensity | 1649.51 | < .001 | 0.91 |
| study x hand | 2.57 | .113 | < 0.001 |
| study x intensity | 0.36 | .545 | 0.002 |
| hand x intensity | < 0.001 | .978 | < 0.001 |
| study x hand x intensity | 0.39 | .528 | < 0.001 |

| Table S5  Behavioral results of the study comparison ANOVA using unpleasantness ratings. | | | |
| --- | --- | --- | --- |
| Effects | *F*_(1,81)_ | *p_(two-tailed)_* | *gen. η^2^* |
| study | 11.37 | .001 | 0.73 |
| hand | 11.78 | < .001 | 0.002 |
| intensity | 576.74 | < .001 | 0.75 |
| study x hand | 3.09 | .083 | < 0.001 |
| study x intensity | 3.42 | .068 | 0.02 |
| hand x intensity | 0.75 | .329 | < 0.001 |
| study x hand x intensity | 1.77 | .188 | < 0.001 |

##### R.4 Region of interest analyses

Below we report the full ANOVA tables of the fMRI results (pooled activation of all ROIs and single ANOVAs for each ROI) as well as the post hoc Bayesian analyses.

| Table S6  ANOVA using pooled activation of seven ROIs. | | | | |
| --- | --- | --- | --- | --- |
| Effects | *df* | *F* | *p_(two-tailed)_* | *gen. η^2^* |
| hand | 1,44 | 0.16 | .689 | < .001 |
| intensity | 1,44 | 13.21 | < .001 | 0.016 |
| roi^S^ | 6,264 | 20.50 | < .001 | 0.144 |
| hand x intensity | 1,44 | 3.09 | .086 | 0.002 |
| hand x roi^S^ | 6,264 | 3.35 | .003 | 0.002 |
| intensity x roi^S^ | 6,264 | 20.19 | < .001 | 0.028 |
| hand x intensity x roi^S^ | 6,264 | 4.61 | < .001 | 0.002 |
| *Note.* Effects marked with an “S” had a significant Mauchly's test for sphericity and corresponding *p*-values are reported using Greenhouse Geisser sphericity correction. The following regions of interest were included: l/rAI = left/right anterior insula, aMCC = anterior midcingulate cortex, l/rS1 = left/right primary somatosensory cortex, l/rS2 = left/right secondary somatosensory cortex. | | | | |

| Table S7  ANOVA of left anterior insula. | | | |
| --- | --- | --- | --- |
| Effects | *F*_(1,44)_ | *p_(two-tailed)_* | *gen. η^2^* |
| hand | 1.21 | .278 | 0.002 |
| intensity | 39.98 | < .001 | 0.162 |
| hand x intensity | 8.93 | .005 | 0.015 |

| Table S8  ANOVA of right anterior insula. | | | |
| --- | --- | --- | --- |
| Effects | *F*_(1,44)_ | *p_(two-tailed)_* | *gen. η^2^* |
| hand | 0.42 | .521 | < .001 |
| intensity | 9.09 | .004 | 0.040 |
| hand x intensity | 6.47 | .015 | 0.012 |

| Table S9  ANOVA of anterior midcingulate cortex. | | | |
| --- | --- | --- | --- |
| Effects | *F*_(1,44)_ | *p_(two-tailed)_* | *gen. η^2^* |
| hand | 0.23 | .636 | < 0.001 |
| intensity | 16.88 | < .001 | 0.053 |
| hand x intensity | 3.64 | .063 | 0.003 |

| Table S10  ANOVA of left primary somatosensory cortex. | | | |
| --- | --- | --- | --- |
| Effects | *F*_(1,44)_ | *p_(two-tailed)_* | *gen. η^2^* |
| hand | 0.03 | .862 | < 0.001 |
| intensity | 3.85 | .056 | 0.003 |
| hand x intensity | 0.39 | .534 | < 0.001 |

| Table S11  ANOVA of right primary somatosensory cortex. | | | |
| --- | --- | --- | --- |
| Effects | *F*_(1,44)_ | *p_(two-tailed)_* | *gen. η^2^* |
| hand | 7.59 | .008 | 0.007 |
| intensity | 1.10 | .300 | 0.002 |
| hand x intensity | < 0.001 | .988 | < 0.001 |

| Table S12  ANOVA of left secondary somatosensory cortex. | | | |
| --- | --- | --- | --- |
| Effects | *F*_(1,44)_ | *p_(two-tailed)_* | *gen. η^2^* |
| hand | 0.26 | .614 | < 0.001 |
| intensity | 0.03 | .853 | < 0.001 |
| hand x intensity | 0.21 | .885 | < 0.001 |

| Table S13  ANOVA of right secondary somatosensory cortex. | | | |
| --- | --- | --- | --- |
| Effects | *F*_(1,44)_ | *p_(two-tailed)_* | *gen. η^2^* |
| hand | 4.00 | .052 | 0.005 |
| intensity | 5.29 | .026 | 0.010 |
| hand x intensity | 0.54 | .468 | < 0.001 |

| Table S14  Overview of the results in the Bayesian t-tests for the seven ROIs. | | | | |
| --- | --- | --- | --- | --- |
| **Region of interest** | **MNI coordinates** | **Prior** | **BF_01_** | **BF_10_** |
| aMCC | -2 23 40 | Cauchy (0, .707) | 0.92 | 1.09 |
| lAI | -40 22 0 | Cauchy (0, .707) | 0.11 | 9.11 |
| rAI | 39 23 -4 | Cauchy (0, .707) | 0.28 | 3.55 |
| lS1 | -39 -30 51 | Cauchy (0, .707) | 9.20 | 0.11 |
| rS1 | 36 -36 48 | Cauchy (0, .707) | 6.29 | 0.16 |
| lS2 | -39 -15 18 | Cauchy (0, .707) | 4.11 | 0.24 |
| rS2 | 39 -15 18 | Cauchy (0, .707) | 3.17 | 0.32 |
| *Note.* MNI coordinates are given as x, y and z; AI = anterior insula; aMCC = anterior midcingulate cortex; r/lS1 = right/left primary somatosensory cortex; r/l S2 = right/left secondary somatosensory cortex; BF_01_ = Bayes Factor evidence for null hypothesis (H_0_ vs. H_1_); BF_10_ = Bayes Factor evidence for alternative hypothesis (H_1_ vs. H_0_); BF_10_ = 1/BF_01_. Bayesian *t*-tests were run in JASP, one-sided for activity_right hand_ > activity_left hand_ (lAI, rAI, aMCC) or activity_right hand_ < activity_left hand_ (lS1, rS1, lS2, rS2). | | | | |
